# Type-3 Secretion System–induced pyroptosis protects Pseudomonas against cell-autonomous immunity

**DOI:** 10.1101/650333

**Authors:** Elif Eren, Rémi Planès, Julien Buyck, Pierre-Jean Bordignon, André Colom, Olivier Cunrath, Roland F. Dreier, José C. Santos, Valérie Duplan-Eche, Emmanuelle Näser, Antonio Peixoto, Dirk Bumann, Céline Cougoule, Agnès Coste, Olivier Neyrolles, Petr Broz, Etienne Meunier

## Abstract

Inflammasome-induced pyroptosis comprises a key cell-autonomous immune process against intracellular bacteria, namely the generation of dying cell structures. These so-called pore-induced intracellular traps (PITs) entrap and weaken intracellular microbes. However, the immune importance of pyroptosis against extracellular pathogens remains unclear. Here, we report that Type-3 secretion system (T3SS)-expressing *Pseudomonas aeruginosa* (*P. aeruginosa*) escaped PIT immunity by inducing a NLRC4 inflammasome-dependent macrophage pyroptosis response in the extracellular environment. To the contrary, phagocytosis of *Salmonella* Typhimurium promoted NLRC4-dependent PIT formation and the subsequent bacterial caging. Remarkably, T3SS-deficient *Pseudomonas* were efficiently sequestered within PIT-dependent caging, which favored exposure to neutrophils. Conversely, both NLRC4 and caspase-11 deficient mice presented increased susceptibility to T3SS-deficient *P. aeruginosa* challenge, but not to T3SS-expressing *P. aeruginosa.* Overall, our results uncovered that *P. aeruginosa* uses its T3SS to overcome inflammasome-triggered pyroptosis, which is primarily effective against intracellular invaders.

**Importance:** Although innate immune components confer host protection against infections, the opportunistic bacterial pathogen *Pseudomonas aeruginosa* (*P. aeruginosa*) exploits the inflammatory reaction to thrive. Specifically the NLRC4 inflammasome, a crucial immune complex, triggers an Interleukin (IL)-1β and -18 deleterious host response to *P. aeruginosa*. Here, we provide evidence that, in addition to IL-1 cytokines, *P. aeruginosa* also exploits the NLRC4 inflammasome-induced pro-inflammatory cell death, namely pyroptosis, to avoid efficient uptake and killing by macrophages. Therefore, our study reveals that pyroptosis-driven immune effectiveness mainly depends on *P. aeruginosa* localization. This paves the way toward our comprehension of the mechanistic requirements for pyroptosis effectiveness upon microbial infections and may initiate targeted approaches in order to ameliorate the innate immune functions to infections.

**Graphical abstract:** Macrophages infected with T3SS-expressing *P. aeruginosa* die in a NLRC4-dependent manner, which allows bacterial escape from PIT-mediated cell-autonomous immunity and neutrophil efferocytosis. However, T3SS-deficient *P. aeruginosa* is detected by both NLRC4 and caspase-11 inflammasomes, which promotes bacterial trapping and subsequent efferocytosis of *P. aeruginosa*-containing-PITs by neutrophils.

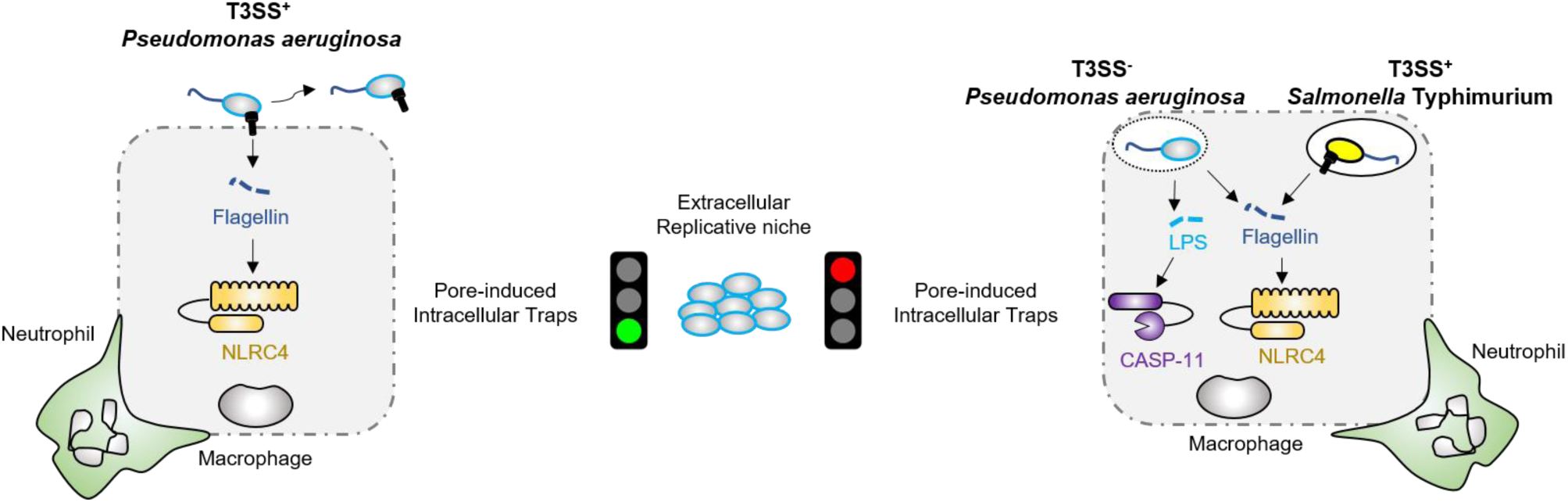

## Introduction

Inflammasomes are pro-inflammatory cytosolic complexes whose activation leads to auto-processing of the protease caspase-1. Caspase-1 activation triggers the cleavage and release of the pro-inflammatory cytokines, interleukin (IL)-1β and IL-18 (1), as well as a pro-inflammatory form of cell death, called pyroptosis. Cleavage and activation of the pore forming effector gasdermin D (GSDMD) by caspase-1 or -11 can induce pyroptosis (2, 3). Several sensors contribute to inflammasome formation, including AIM2-like receptors (AIM2), PYRIN, and a subset of NOD-like receptors (NLRs), namely NLRP1/NLRP1B, NLRP3, NLRP6, NLRP7 and NLRC4, and the caspase-11-induced non-canonical inflammasome pathway (1).

Activation of the NLRC4 inflammasome follows an original path as it requires additional helper NLRs – the neuronal apoptosis inhibitory proteins (NAIPs) – that, upon ligand recognition, form hetero-oligomeric complexes with NLRC4 and promote its activation (4–10). Both NAIP5 and NAIP6 directly recognize cytosolic flagellin while NAIP-1 and -2 detect the type-3 secretion system (T3SS) apparatus needle and rod subunits, respectively (1, 6). Importantly, resulting IL-1β and IL-18 cytokine secretion mediates intracellular bacterial clearance by inducing, respectively, phagocyte recruitment and the production of the microbicidal cytokine interferon (IFN)-γ (1, 6). A recently discovered and understudied cell-autonomous immune process known as pyroptosis promotes intracellular bacteria entrapment and weakening in pore-induced intracellular trap structures (PITs) (11–13), which facilitates subsequent bacterial elimination through efferocytosis (13–16). However, host-adapted bacteria can inhibit PITs formation, which enables bacterial proliferation (14).

While NLRC4 activation confers protection to various pathogens, including *Salmonella* Typhimurium, *Burkholderia pseudomallei* and *Legionella pneumophila* (1, 6), it also drives host susceptibility to the opportunist bacteria *Pseudomonas aeruginosa* (*P. aeruginosa*) (17, 18). In particular, *P. aeruginosa* T3SS and flagellin components induce NLRC4-mediated IL-1β and IL-18 secretion, which inhibits protection mediated by both Th17 cells and anti-microbial peptides produced by airway epithelial cells (17, 18). This apparent paradox underlies complex host-microbe interactions, as NLRC4 confers protection against other pathogens that express T3SS and flagellin. Here, we examined the underlying host-microbe mechanisms through which NLRC4 specifically drives susceptibility to *P. aeruginosa*. We used *Salmonella enterica* serovar Typhimurium (SL1344) and PAO1, a *P. aeruginosa* strain, that expresses a functional T3SS (T3SS^+^) in parallel with an isogenic mutant of this strain, inactivated in the T3SS transcriptional regulator *exs*A, unable to express a functional T3SS apparatus (T3SS^-^).

## Results

### Bacterial localization induces differential pyroptosis-dependent cell-autonomous response in macrophages

To decipher the molecular mechanisms underlying NLRC4-mediated sensing of the bacterial pathogen *P. aeruginosa*, we infected bone marrow-derived macrophages (BMDMs) from wild-type (WT) and *Nlrc4^−/−^* mice with either T3SS-expressing (T3SS^+^) *P. aeruginosa* and *S*. Typhimurium strains (19) and we measured the capacity of both strains to trigger NLRC4 inflammasome-dependent pyroptotic cell death and IL-1β release. In agreement with previous published reports, these data indicate that both bacteria triggered cell-death in a T3SS and NLRC4-dependent manner within 3 hours of infection (MOI 15) (Fig. 1A, Fig. S1A).

**Fig. 1:**
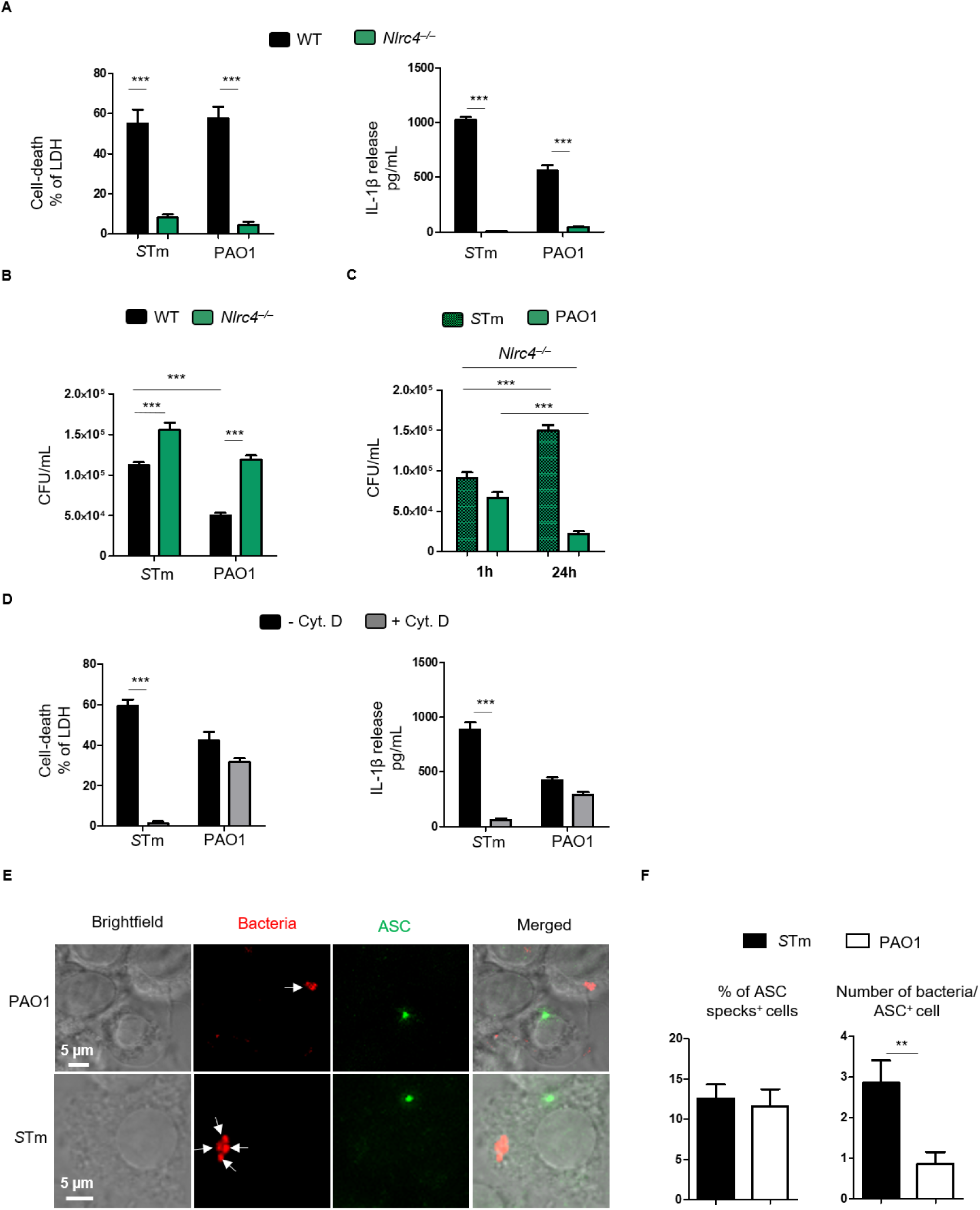
Efficient pyroptosis-induced PIT response in macrophages depends on bacterial localization. BMDMs were primed with 100 ng/mL of the TLR2 ligand Pam_3_CSK_4_ for 16 h to induce pro-IL-1β expression and then infected with *S*. Typhimurium and *P. aeruginosa* for various times. **(A)** Measurement of LDH and IL-1β release in WT and *Nlrc4*^−/−^ BMDMs infected for 3 h with PAO1 and *S*. Typhimurium at an MOI of 15. **(B)** Phagocytosis scoring of WT or *Nlrc4*^−/−^ BMDMs infected for 1 hour with PAO1 and *S*. Typhimurium at an MOI of 15. **(C)** CFU evaluation of *Nlrc4*^−/−^ BMDMs infected for various time (1-24h) with PAO1 and *S*. Typhimurium at an MOI of 15. **(D)** Cell death (LDH) and IL-1β release evaluation in WT BMDMs, pre-incubated or not with cytochalasin D (0.2µg/mL) for 30 minutes, and then infected with either *P. aeruginosa*, *S*. Typhimurium (MOI 15). **(E, F)** Microscopy illustrations and quantifications of PIT (ASC^+^ cells)-associated *P. aeruginosa* and *S*. Typhimurium in WT BMDMs infected for 1H with an MOI 3. Graphs show mean and s.d of quadruplicate wells (A and D) from three independent experiments pooled together. CFUs scoring are representative of one experiment performed 5 times. Regarding microscopy quantification (E, F), 10 fields containing approximately 200 cells were quantified using the Image J software.

While *S.* Typhimurium and *P. aeruginosa* have a different niche tropism (intracellular for *Salmonella* and extracellular for *Pseudomonas*), we hypothesized that the NLRC4 response might lead to different bacterial fate in macrophages. Consequently, CFU assays after 1 hour of infection (MOI 15) found more intracellular *S.* Typhimurium than *P. aeruginosa* in WT BMDMs (Fig. 1B). Strikingly, such differences in intracellular numbers of *P. aeruginosa* was partially lost in *Nlrc4^−/−^* macrophages and independent from *P. aeruginosa* T3SS-injected exoenzymes STY (Fig. 1B, Fig. S1B), suggesting that the NLRC4 inflammasome response contributed to *P. aeruginosa* avoiding macrophage uptake. Whereas *Salmonella* could efficiently establish an intracellular replicative niche in *Nlrc4*^-/-^ BMDMs, intracellular *P. aeruginosa* failed to do so after 24 hours of infection (Fig. 1C), which suggests that *P. aeruginosa* exploits NLRC4-dependent response to avoid macrophage intracellular-autonomous immunity. Phagocytosis is the main process by which macrophages engulf bacteria. Hence, inhibiting phagocytosis by cytochalasin D reduced NLRC4-dependent cell death and IL-1β release upon infection with *S*. Typhimurium but not with *P. aeruginosa* (Fig. 1D). Remarkably, T3SS^+^-induced phagocytosis-independent activation of the NLRC4 inflammasome was specific to only *P. aeruginosa*, as other T3SS-expressing bacteria triggered NLRC4 inflammasome responses in a phagocytosis-dependent manner (Fig. S1C). To rule out that actin polymerization might directly control NLRC4 activation, we electroporated purified flagellin in WT and *Nlrc4*^-/-^ BMDMs in presence or absence of cytochalasin D. Cell death evaluation showed that cytochalasin D did not modify cytosolic flagellin-induced NLRC4 inflammasome response (Fig. S1D). Thus, these results identified the unique capability of T3SS-expressing *P. aeruginosa* to promote NLRC4-inflammasome activation in a phagocytosis-independent manner.

“Pore-induced Intracellular Traps” (PITs) entrap intracellular *S*. Typhimurium (11–13). As *P.aeruginosa* induced phagocytosis-independent activation of the NLRC4 inflammasome, we reasoned that such process might be a virulence strategy to escape bacterial caging into PITs. Therefore, we infected primary WT BMDMs with both *P. aeruginosa* and *S*. Typhimurium strains (MOI 3) for 1h to induce PIT formation and evaluated the number of bacteria associated with pyroptotic structures (e.g. ASC^+^ cells). Strikingly, our results found higher amount of *Salmonella* associated within PIT structures than *P. aeruginosa*, showing that *P. aeruginosa*-activated NLRC4 inflammasome allowed escape from PIT-driven intracellular trapping (Fig. 1E, F).

Then, we aimed to determine whether pyroptosis have a differential *in vivo* regulatory function after *P. aeruginosa* and *S*. Typhimurium exposure. Using a peritoneal mouse model of infection (3×10^6^ CFUs, 6h) with either strains, we evaluated the contribution of pyroptosis on the early clearance of each bacterial strain in WT or in *gasderminD^-/-^* (*gsdmD ^-/-^*) mice, which are deficient at inducing pyroptosis. Therefore, mice lacking *gsdmD* showed an early (6h) susceptibility to *S*Tm challenge whereas they presented improved resistance to *P. aeruginosa* infection (Fig. S1F). Overall, these results demonstrated that *P. aeruginosa* exploited the NLRC4 inflammasome to escape pyroptosis-mediated capture and sequestration, a process that was reversed in absence of *Nlrc4*.

### *Pseudomonas* triggers T3SS-independent NLRC4 and Caspase11 inflammasome activation in macrophages

Our results showed that both *S.* Typhimurium and *P. aeruginosa*-induced differential PIT-dependent bacterial trapping. Yet, we wanted to rule out that the observed phenotype could be driven by intrinsic properties of each bacterial strain, such as flagellin and T3SS immunogenicity and/or expression levels.

Therefore, we relied on a surprising observation where we found that infection of macrophages with a T3SS-deficient strain of *P. aeruginosa* still triggered early (3 hours) residual NLRC4 response and, late (4-10 hours) caspase11 non-canonical inflammasome pathway (Fig. 2A; Figs. S2A, B). Even though high doses (MOI 25, 50 or 100) of T3SS^-^ *P. aeruginosa* were necessary to induce reduced levels of NLRC4 activation, residual inflammasome response was both still NLRC4- and -flagellin dependent, as T3SS^-^/*fliC*^-^ strains failed to elicit measurable IL-1β release as well as caspase-1 and gasdermin-D (GSDMD) processing (Figs. 2A, B; Fig. S2C). Furthermore, we noted that flagellin complementation in both T3SS^+^ and T3SS^-^ strains restored NLRC4-driven cell death and IL-1β release (Figs. S2D, E). However, only live T3SS-deficient *P. aeruginosa* induced NLRC4-dependent IL-1β release as it was not detected using heat-killed (HK) and PFA-killed bacteria (20) (Fig. S2F). To exclude that genetic inactivation of *exsA* allowed residual T3SS expression, we tested *P. aeruginosa* lacking PscC, a key T3SS structural component (21). Importantly, *pscC* and *exsA*-deficient bacteria triggered comparable levels of NLRC4-dependent cell death and IL-1β release (Fig. S2G). These results indicate that activation of the NLRC4 inflammasome was not caused by residual T3SS expression.

**Fig. 2:**
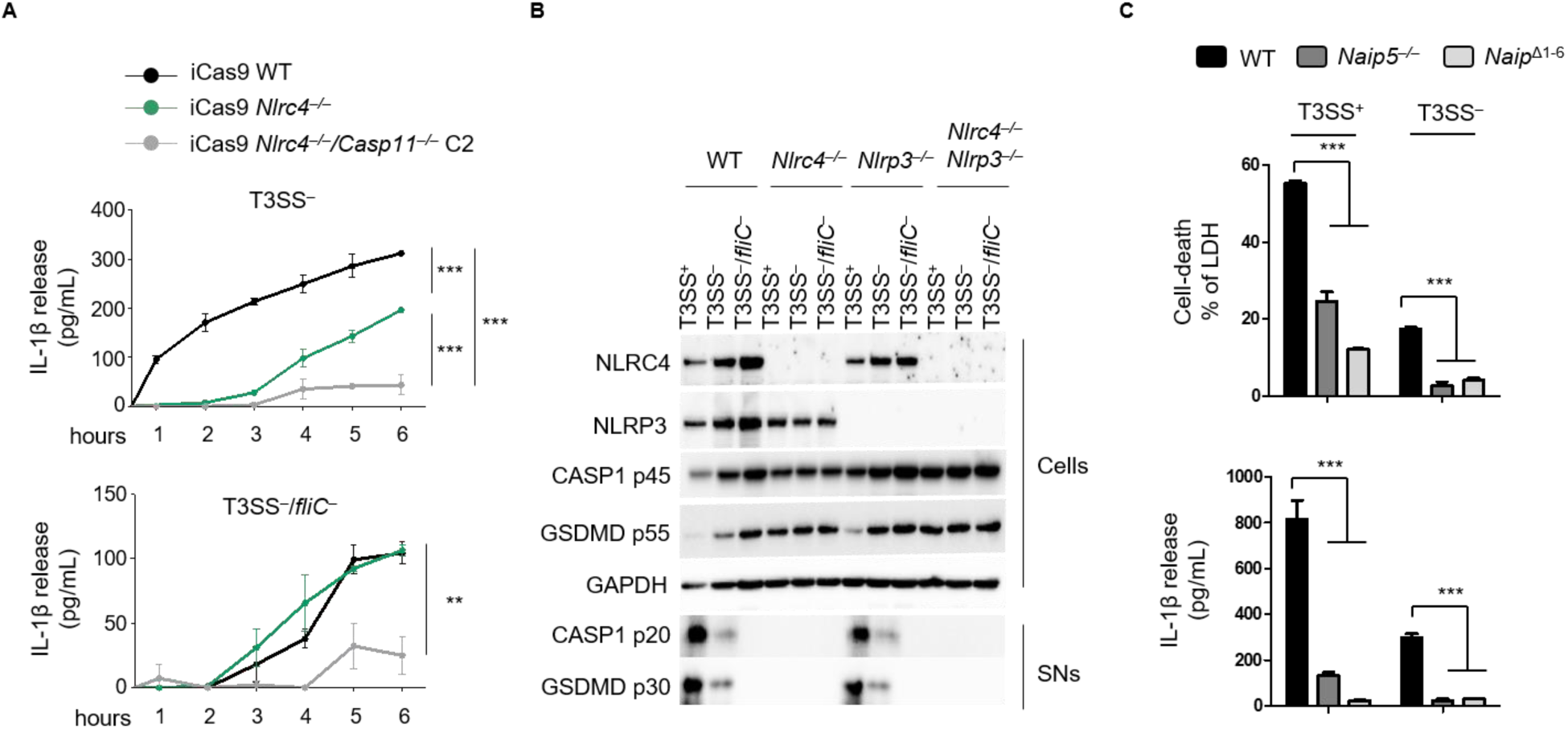
T3SS^-^ *P. aeruginosa* activate both NLRC4 and Caspase11 inflammasomes. PAM3CSK4-primed (100 ng/mL) BMDMs were infected for various times with bacterial strains PAO1 T3SS^+^, T3SS^-^ or T3SS^-^/*fliC*^-^ at an MOI of 25, unless otherwise stated. **(A)** Kinetics of IL-1β released by immortalized WT, *Nlrc4*^−/−^ or *Nlrc4*^−/−^/*Casp11*^−/−^ BMDMs infected with PAO1 T3SS^-^ or T3SS^-^/*fliC*^-^. **(B)** Western blot examination of processed caspase-1 (p20) and gasdermin-D (p30) in supernatants and pro-caspase-1 (p45), pro-gasdermin-D (p55), NLRP3, NLRC4 and GAPDH in cell lysates of WT, *Nlrc4^−/−^*, *Nlrp3^−/−^* and *Nlrc4^−/−^/Nlrp3^−/−^* BMDMs infected for 3 h with PAO1 T3SS^+^, T3SS^-^ or T3SS^-^/*fliC*^-^. **(C)** Measurement of LDH and IL-1β release from WT, *Naip5*^−/−^ or *Naip*^Δ1-6^ BMDMs infected for 3 h with PAO1 T3SS^+^ or T3SS^-^. Graphs show mean and s.d of quadruplicate wells **(A, C)** pooled from three independent experiments. Immunoblotting **(B)** is representative of one experiment performed two times.

In the host, cytosolic NAIP proteins bind flagellin or T3SS components and initiate NLRC4 oligomerization (6). To confirm direct involvement of flagellin in NLRC4 activation, we infected macrophages lacking either *Naip5* (*Naip5^−/−^*), the main flagellin sensor, or the 5 different *Naips* (*Naip*^Δ1-6^), with either T3SS^+^ or T3SS^-^ strain. Both *Naip5^−/−^* and *Naip*^Δ1-6^ BMDMs failed to undergo pyroptosis and release IL-1β when infected with either strain (Fig. 2C). Based on these results, we conclude that flagellin directly triggered the NLRC4 inflammasome response during T3SS^-^ *P. aeruginosa* infection.

Altogether, these findings reveal, in addition to the Caspase-11 non canonical inflamamsome, an unrecognized and unexpected capacity of *P. aeruginosa* to alternatively activate the NAIP5/NLRC4 inflammasome in a T3SS-independent yet flagellin-dependent manner. These findings further suggest that flagellin can reach the host cell cytosol independently of bacterial secretion systems.

### T3SS^-^ *P. aeruginosa*-containing damaged phagosomes associate to sequential activation of NLRC4 inflammasome and Caspase-11

While our results indicated that flagellin and LPS reache the host cytosol in absence of a functional T3SS, the underlying mechanism still remained unknown. We hypothesized that a key candidate involves direct entry of *P. aeruginosa* into the intracellular compartment. Therefore we evaluated whether T3SS-independent activation of both NAIP5/NLRC4 and Caspase-11 inflammasomes required *P. aeruginosa* phagocytosis. Consequently, infection of macrophages with T3SS^-^ *P. aeruginosa* showed that phagocytosis inhibition abrogated cell death, IL-1β release and Caspase-1 processing (Fig. 3A-C). Hence, these results show that, in absence of a functional T3SS, *P. aeruginosa* induces inflammasome response in a phagocytosis-dependent manner. As phagocytosis of T3SS-deficient *P. aeruginosa* is a prerequisite for inflammasome response, we hypothesized that NAIP5/NLRC4 and Caspase11 responses required that both flagellin and LPS reached the cytosol by leaking from *P. aeruginosa*-containing phagosomes. To visualize phagosomal membrane alterations, we probed for galectin-3 (GAL3), a lectin that binds galactosides on permeabilized host cell endovesicles (22). If *P. aeruginosa*-containing phagosomes were compromised, we expected GAL3 recruitment around bacteria. To avoid unwanted GAL3 recruitment to permeabilized phagosomes from pyroptosis and not *P. aeruginosa* specifically, we infected immortalized *Casp1^−/−^/11^−/−^* macrophages with two T3SS mutants (e.g. T3SS^-^ and T3SS^-^/*fliC*^-^) for 3 hours. Confocal microscopy revealed that GAL3 did target a small proportion (∼ 6-10%) of intracellular T3SS^-^ and T3SS^-^/*fliC*^-^ bacteria (Fig. 3D). We then reasoned that cells with active inflammasomes, *i.e.* containing ASC specks, should also contain at least one GAL3-positive bacterium. Using macrophages expressing an active inflammasome (ASC specks^+^), we quantified the percentage of cells that had at least one GAL3-positive *P. aeruginosa*. We consistently found that approximatively 45% of i*Casp-1*^−/−^*/11*^−/−^ macrophages presenting an ASC speck were also positive for GAL3-stained T3SS^-^ *P. aeruginosa* (Figs. 3D, E), but we did not detect any specks using T3SS^-^/*fliC*^-^ strain (Fig. 3E). *Casp1^−/−^/11^−/−^* deficient BMDMs can form a NLRC4 inflammasome that recruit caspase-8 that also triggers cell death. To verify that in *Casp1^−/−^/11^−/−^* iBMDMs, GAL3 recruitment to intracellular bacteria did not require caspase-8, we removed *Nlrc4* in GAL3-mcherry expressing *Casp1^−/−^/11^−/−^* iBMDMs (Fig. S3A) and quantified GAL3 recruitment to T3SS^-^ *P. aeruginosa*. We did not detect any defect for bacteria targeted by GAL3-mecherry in the *Nlrc4*^-/-^/*Casp1^−/−^/11^−/−^* iBMDMs, hence confirming that T3SS- *P. aeruginosa* is targeted by GAL3 independently of the NLRC4-dependent pathway (Fig. S3B). Overall, these results show that T3SS^-^ *P. aeruginosa* trigger NLRC4 inflammasome from altered phagosomes.

**Fig. 3:**
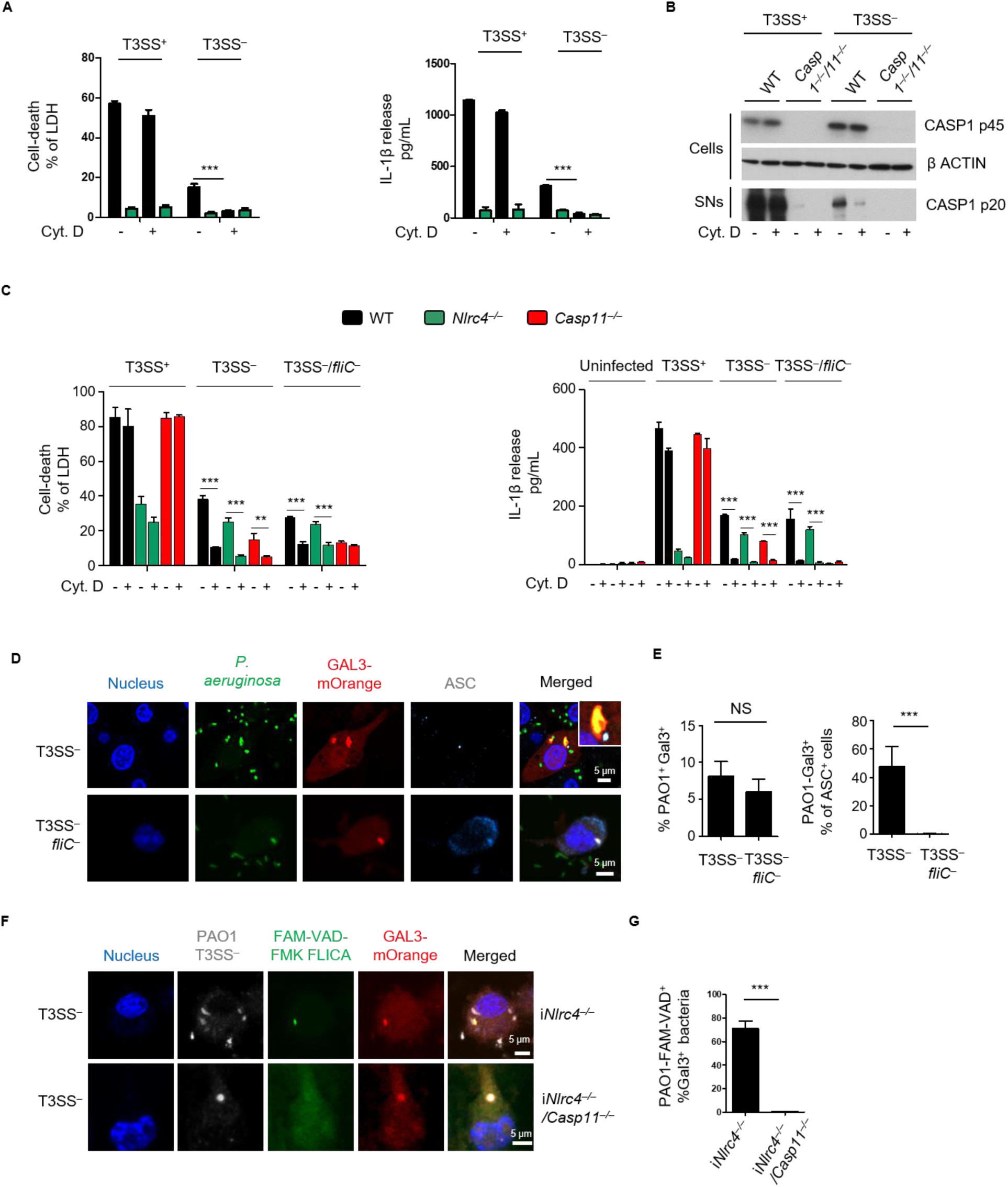
T3SS^-^ *P. aeruginosa* associate to damaged phagosomes to promote inflammasome response. PAM3CSK4-primed BMDMs were infected with various *Pseudomonas aeruginosa* strains at an MOI of 25, unless specified. **(A)** LDH and IL-1β released by WT and *Nlrc4^−/−^* BMDMs, pre-incubated or not with cytochalasin D (0.2µg/mL) for 30 minutes, and then infected for 3 h with PAO1 T3SS^+^ or T3SS^-^. **(B)** Western blot analysis of cleaved CASP1 (p20) in cell supernatants and of pro-CASP1 and β-actin in cell extracts of WT and *Casp1^−/−^/11^−/−^* BMDMs infected for 3 h with T3SS^+^ or T3SS^-^ PAO1. **(C)** LDH and IL-1β release in WT, *Nlrc4*^−/−^ or *Casp11*^−/−^ BMDMs, pre-incubated or not with cytochalasin D (0.2µg/mL) for 30 minutes, and then infected for 10 h with PAO1 T3SS^+^, **(D, E)** Fluorescence microscopy observations and quantifications of galectin-3 targeted PAO1 T3SS^-^ or T3SS^-^/*fliC*^-^ (MOI 25, 3 hours infection) in *Casp-1^−/−^/11^−/−^* immortalized BMDMs transduced with galectin-3-morange. **(F, G)** Observations and quantifications of FAM-VAD-FMK FLICA probe recruitment on GAL-3-mOrange^+^ *Pseudomonas* aeruginosa (T3SS^-^) in IFNγ-primed immortalized (i) i*Nlrc4*^−/−^/GAL3-morange and i*Nlrc4*^−/−^/*Casp11*^−/−^/GAL3-mOrange BMDMs 3 h after infection. Caspase-11 is in green, galectin-3 in red, bacteria in grey and nuclei in blue. Graphs show mean and s.d. **(A, C)** represent pooled data from at least three independent experiments performed in quadruplicate. **(B)** is representive of one experiment performed in duplicate. **(D, F)** Microscopy quantification of 10 fields containing approximately 200 cells performed using the Image J software.

Given that T3SS^-^ *P. aeruginosa* were exposed to the cytosol, we speculated that phagosome alterations could also expose *P. aeruginosa* LPS to the host cell cytosol, which might promote caspase11 recruitment and activation. Thurston et al. showed that caspase11 was recruited to cytosolic Salmonella in infected epithelial cells (25). So, we sought to determine whether caspase11 associated with compromised *P. aeruginosa*-containing phagosomes. Microscopy observations of galectin-3+ (GAL3+) bacteria in IFNγ-primed *Nlrc4*^−/−^ iBMDMs showed recruitment of both GAL3 and active caspase on T3SS^-^ *P. aeruginosa* after 3 hours of infection, which was not observed in *Nlrc4*^−/−^/*Casp11*^−/−^ CRISPR-Cas9 iBMDMs (Fig. 3F). These results confirm that bacteria accessible to the cell cytosol directly recruited caspase-11.

Altogether, these results indicate that both NLRC4 and Caspase11 directly detected intracellular T3SS^-^ *P. aeruginosa* flagellin and LPS from compromised phagosomes.

### Pyroptosis-induced PITs is efficient only against T3SS-deficient *Pseudomonas*

As we previously showed that T3SS-expressing *P. aeruginosa* and *S*. Typhimurium triggered differential PIT responses in macrophages, we evaluated whether pyroptosis-induced PITs also promoted restriction of intracellular *P. aeruginosa*. Primary WT BMDMs were infected with T3SS^+^ or T3SS^-^ for 3 h to induce PIT formation. Time-lapse microscopy revealed that cells with compromised plasma membranes (e.g. positive for the DNA impermeant binding probe TO-PRO-3) contained a high number of intracellular T3SS^-^ *P. aeruginosa*, while few T3SS^+^ *P. aeruginosa* were associated with dead cells (**Movie 1 that refers to T3SS^+^ PAO1-infected cells, Movie 2 that refers to T3SS^-^PAO1-infected cells**). In addition, confocal microscopy experiments showed that PITs (*i.e.* ASC^+^ cells) were enriched specifically with T3SS^-^ bacteria (Fig. 4A, B). Based on these results, we conclude that T3SS-expressing *P. aeruginosa* hijacked PIT-mediated intracellular bacterial entrapment and weakening, a process reversed in the absence of T3SS expression.

**Fig. 4:**
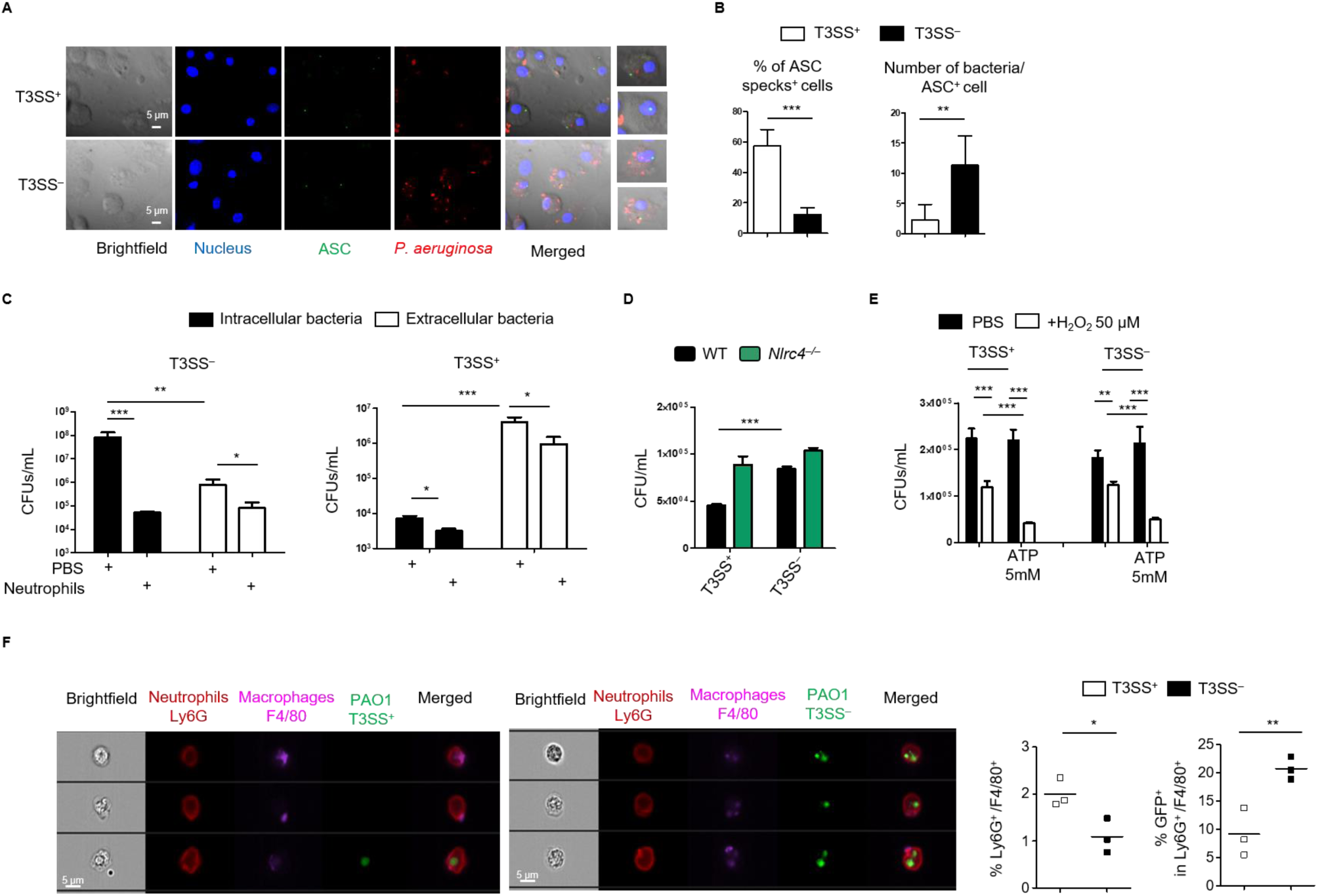
Pyroptosis-induced *P. aeruginosa* entrapment and elimination is mostly efficient against T3SS-deficient bacteria. Unprimed BMDMs were infected with various *Pseudomonas aeruginosa* strains at an MOI of 25, unless otherwise specified. **(A,B)** Microscopy illustrations and quantifications of PIT (ASC^+^ cells)-associated *P. aeruginosa* T3SS^+^ or T3SS^-^ in WT BMDMs infected for 3h. **(C)** Bacterial weakness induced by pyroptosis evaluated by LB agar plating of intracellular and extracellular PAO1 T3SS^+^ and T3SS^-^ from WT infected BMDMs after PBS or neutrophil exposure (1 h 30). **(D)** CFU scoring of WT or *Nlrc4*^−/−^ BMDMs infected for 1 hour with PAO1 T3SS^+^ or T3SS^-^ at an MOI of 25. **(E)** Bacterial weakness induced by pyroptosis (ATP, 5 mM, 2 h) evaluated by LB agar plating of intracellular *P. aeruginosa* T3SS^+^ and T3SS^-^ (MOI 25, 1h) from *Nlrc4*^−/−^ infected BMDMs after PBS or H_2_O_2_ exposure (50 µM). **(F)** ImagestreamX quantification of the % of (i) Neutrophils (Ly6G^+^)/macrophage (F4/80^+^) and, (ii) Neutrophils (Ly6G^+^)/macrophages (F4/80^+^)/bacteria (GFP^+^) in the peritoneal cavity of WT mice infected for 4 h with 3.10^6^ CFUs of T3SS^+^ or T3SS^-^ *P. aeruginosa*. Single dots are representative of each individual mouse infected with either strain of *P. aeruginosa*. Images represent the acquisition of more than 100 000 total events. Upper image panel and lower image panel show representatives images from the Neutrophils (Ly6G+)/macrophages (F4/80+) gate obtained from mice infected with T3SS^+^ or T3SS^-^ *P. aeruginosa* respectively. Here, three independent experiments were conducted. All data **(A-F)** are expressed as mean and s.d. Data are representative of three independent experiments. **Movie 1** shows T3SS^+^ PAO1 mcherry entrapment into dead cells and refers to Fig. 4. **Movie 2** shows T3SS^-^ PAO1 mcherry entrapment into dead cells and refers to Fig. 4.

The capability of PITs to entrap mostly T3SS-deficient bacteria suggested that these structures might also have a direct or an indirect microbicidal function against these strains, as suggested by *S*. Typhimurium infection (13, 15). We then infected primary WT BMDMs with either T3SS^+^ or T3SS^-^ *P. aeruginosa* for 3h and monitored the microbicidal potential of PITs. PITs-associated or –unbound bacteria were harvested and exposed to purified neutrophils or PBS for 1h30. We quantified CFUs, which showed that PITs-associated bacteria specifically were more susceptible to neutrophil killing than the PITs-unbound fraction (Fig. 4C). Since phagocytosis of both T3SS^+^ and T3SS^-^ *P. aeruginosa* was comparable in *Nlrc4* deficient BMDMs (Fig. 4D), we hypothesized that pyroptosis induction in these cells would weaken both bacteria to the same extent. We infected *Nlrc4*^-/-^ BMDMs with both T3SS^+^ and T3SS^-^ *P. aeruginosa* strains and subjected them to ATP stimulation to ensure similar NLRP3 inflammasome-dependent pyroptosis induction in both conditions. Intracellular bacteria were harvested, counted and exposed to a secondary stress signal (i.e. H_2_O_2_). Both intracellular *P. aeruginosa* strains showed the same increased susceptibility to H_2_O_2_ (Fig. 4E), thus demonstrating that PIT-driven intracellular trapping was only efficient against T3SS-deficient *P. aeruginosa*. Finally, we performed a peritoneal mouse model of infection (3×10^6^ CFUs, 4h) with GFP-expressing T3SS^+^ or T3SS^-^ *P. aeruginosa*, and evaluated bacteria-containing dead macrophages efferocytosis by neutrophils (13). Strickingly, recruited neutrophils (Ly6G^+^) having efferocytosed bacteria-containing macrophages (Ly6G^+^/F4/80^+^/GFP^+^) was more efficient against T3SS^-^ *P. aeruginosa* than to T3SS^+^ bacteria (Fig. 4E; Fig. S4A). Overall, these results demonstrated that *P. aeruginosa* used its T3SS to escape PIT-mediated capture and sequestration. This process was reversed in absence of T3SS-expression or of *Nlrc4*.

### Both NLRC4 and Caspase-11 specifically protect against acute infection with T3SS-deficient *P. aeruginosa*

To evaluate the *in vivo* relevance of our findings, we infected WT and *Nlrc4^−/−^* mice with either T3SS^+^ or T3SS^-^ *P. aeruginosa*. Because T3SS^-^ *P. aeruginosa* is highly susceptible to immune defenses (23), we infected mice with a higher dose of this strain. When infected for 18h with T3SS^+^ *P. aeruginosa*, *Nlrc4^−/−^* mice presented lowered bacterial loads in the bronchoalveolar lavage fluid (BALF) than their WT counterparts (Fig. 5A), consistent with a prior report (18). Remarkably, *Nlrc4^−/−^* mice infected with the T3SS^-^ strain showed an increased BALF bacterial loads compared to WT controls after 18h of infection (Fig. 5A). The use of T3SS^-^/*fliC*^-^ *P. aeruginosa* did not show any involvement of NLRC4 on bacterial loads in BALFs, confirming that flagellin was the principal component that mediates *in vivo* NLRC4 response to T3SS-deficient *P. aeruginosa* (Fig. S5A).

**Fig. 5:**
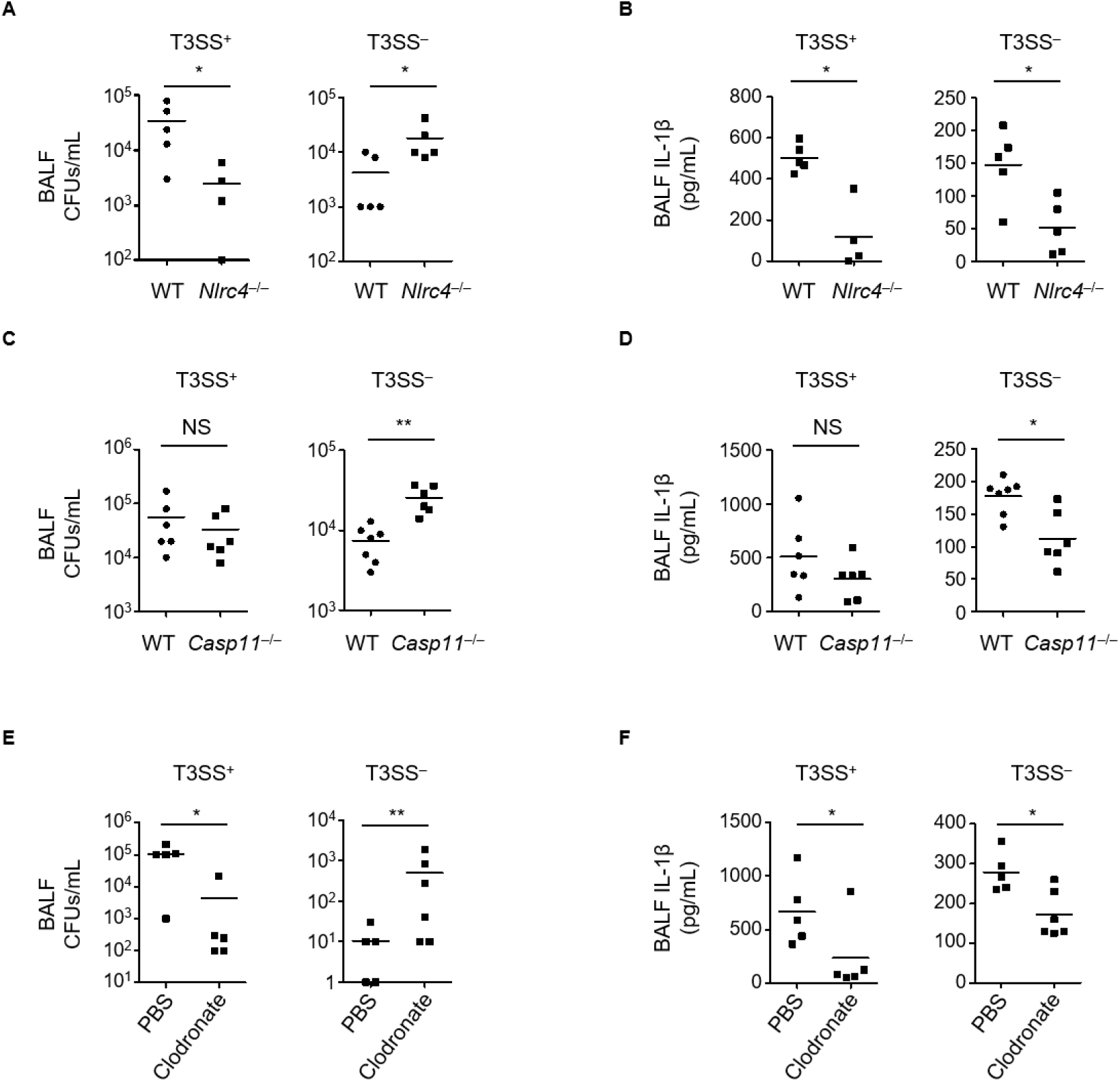
Both NLRC4 and Caspase-11 only protect against acute infection with T3SS-deficient *P. aeruginosa*. **(A, C)** BALF PAO1 CFU scoring after 18 h of WT, *Nlrc4*^−/−^ **(A)** or *Casp11*^−/−^ **(C)** mice infected with either T3SS^+^ or T3SS^-^ *P. aeruginosa* strains at 5×10^6^ (T3SS^+^) and 1.5×10^7^ (T3SS^-^) CFUs respectively. **(B, D)** BALF IL-1β release assay in mice infected as described in **(A, C)**. **(E, F)** Role of alveolar macrophages on the immune response to PAO1. **(E)** PAO1 CFUs scoring in BALFs after 18 h of infection with either the T3SS^+^ or T3SS^-^ *P. aeruginosa* strains in control or clodronate treated mice as in **(A, C)**. **(F)** IL-1β release in BALFs of control or clodronate treated mice infected with either the PAO1 T3SS^+^ or T3SS^-^ strains as in **(B, D)**. All data are representative results of 2 **(C-F)** and 3 **(A, B)** independent experiments.

While strongly reduced, the residual IL-1β levels found in BALFs of mice infected with the T3SS-deficient *P. aeruginosa* strain remained partially dependent on the NLRC4 inflammasome (Fig. 5B), whereas infection with T3SS^-^/*fliC*^-^ bacteria showed no NLRC4-dependent IL-1β release (Fig. S5B). Since we previously showed that caspase-11 could also detect intracellular *P. aeruginosa*, we speculated about the potential immune importance of this pathway *in vivo*. Although *Casp11* deficient mice did not show significant involvement of caspase-11 at controlling the T3SS^+^ strain, T3SS^-^ and T3SS^-^/*fliC*^-^ bacterial infection induced a higher susceptibility in these mice than their WT counterparts (Figs. 5C, D; Figs. S5C, SD). This result verified that both NLRC4 and caspase-11 play paired protective roles only against T3SS-deficient *P. aeruginosa*.

After airway infection, *P. aeruginosa* will encounter alveolar macrophages as the first phagocytic cells. So, we evaluated the importance of alveolar macrophages on inflammasome-triggered differential host responses to *P. aeruginosa*. Mice infected with T3SS^+^ and T3SS^-^ *P. aeruginosa* showed that alveolar phagocyte depletion (Fig. 5E) increased alveolar loads of T3SS^-^ *P. aeruginosa* (Fig. S5E) and strongly impaired IL-1β release in the alveoli (Fig. 5F). These results demonstrated that alveolar macrophages controlled inflammasome-mediated immune responses to *P. aeruginosa* in mice.

Overall, these results demonstrate that T3SS-expressing *P. aeruginosa* exploited the NLRC4-dependent response to their own advantage, but T3SS deficiency uncovered host protective NLRC4- and caspase-11 responses.

## Discussion

Although the NAIP-NLRC4 inflammasome primarily controls intracellularly adapted bacteria *Salmonella* or *Legionella* spp, several studies indicate the critical function of the T3SS-flagellin complex in mediating NLRC4-dependent host susceptibility to *P. aeruginosa* infection is still discussed (17, 18, 24)(25). Regarding *P. aeruginosa* infection, the deleterious functions of both IL-1β and IL-18 on immune control of T3SS-expressing *P. aeruginosa* infection is well documented (18, 26), yet the role of pyroptosis in this process remains unclear.

Here, we provide evidence that NLRC4-dependent pyroptosis of macrophages drive differential outcomes of T3SS-expressing *P. aeruginosa* and *S*. Typhimurium. Whereas *S*. Typhimurium remains entrapped into PITs, as previously demonstrated (13, 16), *P. aeruginosa*-induced extracellular activation of the NLRC4 inflammasome allows bacterial escape from macrophage and pyroptosis-driven cell-autonomous immunity. To our knowledge, such response is unique to *P. aeruginosa*, as other T3SS-expressing bacteria also required uptake to activate the NLRC4 inflammasome. A recent study found that *P. aeruginosa* T3SS triggered a Caspase-3/-7 deleterious host response in an ASC-dependent manner (27). Therefore, it is tempting to speculate that a similar mechanism underlying T3SS-expressing *P. aeruginosa*-induced cell apoptosis could also mediate bacterial escape from host immune responses.

In addition, we uncovered that a T3SS-deficient strain of *P. aeruginosa* also triggered residual, but of immune importance, NAIP5/NLRC4 inflammasome, in a T3SS-independent manner, which is in agreement with findings of Faure et al., where lethal challenge of T3SS-deficient *P. aeruginosa* still induce a NLRC4-dependent response in mice. In addition, T3SS-expressing *P. aeruginosa* triggered inflammasome assembly through a process that did not heavily utilize phagocytosis (28–30). We do not believe that the presence of a genetically encoded phagosomal permeabilization system in *P. aeruginosa* is likely, as only a minority (∼6-10%) of bacteria were accessible to cytosolic galectin-3. Another explanation could involve phagosome maturation-induced *P. aeruginosa* local production of outer-membrane vesicles (OMVs), which could expose bacterial ligands to host cell cytosol sensors (31–34). Conversely, *Nlrc4* deficiency or long-term infection of macrophages promoted a switch in the NLRC4 response to a LPS-induced caspase-11 non-canonical inflammasome path. Although previous results from others and our group indicated that OMVs from *P. aeruginosa* induced non-canonical inflammasome pathway in a phagocytosis-independent, yet endocytosis-dependent manner, our results here demonstrated that T3SS-deficient *P. aeruginosa*-triggered caspase-11 non-canonical inflammasome responses required bacterial uptake (31–34). Future investigations will be critical to determine the specific roles of OMVs and intracellular *P. aeruginosa* for nucleation of the non-canonical inflammasome route. Immunodeficiency, burn-, nosocomial- and ventilator-associated injuries render patients susceptible to chronic *P. aeruginosa* infection, in which most strains are deficient for T3SS and/or flagellin expression (35, 36). A recent report suggested that T3SS^-^ *P. aeruginosa* promoted the non-canonical inflammasome pathway in an IFN- and GBP-dependent manner (37). Consistent with these data, we also showed that caspase-11 protected mice from acute infection with lethal doses of T3SS-deficient *P. aeruginosa*.

The *in vivo* function of NLRC4 to *P. aeruginosa* infection remains unresolved as it can be either protective or deleterious to the host in mouse models (24, 38). Our mouse model of acute infection revealed a deleterious role of NLRC4 on host defenses during exposure to T3SS-expressing *P. aeruginosa* (17, 18). Importantly, we demonstrated that *P. aeruginosa* T3SS deficiency induced both NLRC4- and caspase-11-driven host protection during infection. Attree’s group analyzed isolated naturally T3SS deficient clinical isolates from patients (39). Some strains expressed flagellin and were motile (40). These findings warrant investigating whether these bacteria also induce a NLRC4 dependent response.

Overall, our study uncovered that T3SS-expressing *P. aeruginosa* remained extracellular by triggering cell pyroptosis, a process that enabled bacteria to overwhelm macrophage PITs-induced host immunity. We further demonstrated that phagocytosis of *P. aeruginosa* that lacked T3SS expression triggered host protective pyroptosis in a T3SS-dependent manner through both NLRC4 and caspase-11 inflammasomes. In summary, we propose that bacterial localization drives the effectiveness of the inflammasome response during bacterial infection **(Graphical abstract)**. Future studies will determine if this is a conserved response among bacterial pathogens.

## Methods

All reagents and biological samples used in this study are listed in the table S1

### Mice

*Casp11*^−/−^, *Casp1*^−/−^*/Casp11*^−/−^, *Nlrc4*^−/−^, *Nlrp3^−/−^* and *Gsdmd*^−/−^ mice have been previously described (41–43). Mice were bred in the animal facilities of the University of Basel (Basel, Switzerland) or at the IPBS institute (Toulouse, France). Bones from *Naip5^−/−^* and *Naip^Δ1-6^* mice were a kind gift from R. E Vance (UC Berkeley, USA) (43). Janvier and Charles Rivers companies provided WT mice.

### Animal infections

6-8 age and sex-matched animals (8–10 weeks old) per group were infected intranasally with 5×10^6^ (PAO1 T3SS^+^) or 1.5×10^7^ (PAO1 T3SS^-^ or T3SS^-^/*fliC*^-^) CFUs of mid-late exponential phase *Pseudomonas aeruginosa* in 40µl PBS. Animals were sacrificed 18h after infection and bronchoalveolar fluids (BALFs) were collected in PBS. When specified, cellular contents (flow cytometry), bacterial loads (serial dilutions and CFU plating) and cytokine levels (ELISA) were evaluated. No randomization or blinding were done.

### Clodronate-induced alveolar phagocyte depletion

C57BL/6 mice received intranasal instillation of 40µL of either clodronate or PBS loaded liposomes to deplete alveolar macrophages (18). 48 h later, mice were intranasally infected with either 5×10^6^ (PAO1 T3SS^+^) or 1.5×10^7^ (PAO1 T3SS^-^) CFUs of mid-late exponential phase *Pseudomonas aeruginosa* (OD of 1.0-1.6) in 40µl PBS. 18 h after infection, BALFs were collected and bacterial loads (serial dilutions and CFU plating), cytokine levels (ELISA) and immune cell contents were evaluated. Briefly, cells were pelleted (1000 rpm, 5 minutes) and alveolar macrophages were subsequently stained with a cocktail of fluorochrome-conjugated antibodies detailed in the material section. Cells were then fixed in 4% PFA before fluorescence associated cell sorting (FACS) analysis using a LSRII instrument (BD Biosciences). Data analysis and processing were performed using FlowJo.10 software. The plot FSC-A vs FSC-H was used to discriminate doublets from single cells. Alveolar macrophages were defined as CD11c^+^/F480^+^ in Ly6C^-^/Ly6G^-^/CD19^-^/TCRβ^-^ population.

### Genetic invalidation of *Nlrc4* and *Caspase11* genes in i*Casp1*^−/−^/*Casp11*^−/−^ and i*Nlrc4*^−/−^ immortalized BMDMs

*Nlrc4* was knocked-out using the crispr/cas9 system in *iCasp1*^-/-^/*Casp11*^-/-^ macrophages and *Casp11* was knocked-out from i*Nlrc4*^-/-^ macrophages. Single guide RNAs (sgRNA) specifically targeting *Nlrc4* exon3 (Forward: 5’CACCGTTACTGTGAGCCCTTGGAGC3’ reverse: 5’AAACGCTCCAAGGGCTCACAGTAA-C3’) and *Caspase-11* exon 2 forward (5‘CACCGCTTAAGGTGTTGGAACAGCT3’) reverse (5’AAACAGCTGTTCCAACACCTTAAGC3’) were designed using Benchling tool (Benchling.com), and oligonucleotides were synthesized by Sigma-Aldrich. The gene-specific crispr guide RNA oligonucleotides were hybridized and cloned in Lenti-gRNA-Puromycin vector using BsmBI restriction sites (lentiGuide-Puro, Addgene 52963, from Feng Zhang lab). These constructs were transfected (using lipofectamine 2000) into HEK293T cells together with the lentiviral packaging vector p8.91 (from Didier Trono lab, EPFL, Switzerland) and a envelop VSVg-encoding vector (pMD.2G, Addgene 12259, from Didier Trono lab) for 48 h. Then, viral supernatants were harvested, filtered on 0.45 µm filter. Cas9-expressing recipient-cells either i*Casp1*^−/−^/*Casp11*^−/−^ or i*Nlrc4*^−/−^ (1,000,000 cells/well in 6-well plates) were generated by lentiviral transduction with Cas9-expressing lentiviral vector (lentiCas9-Blast, Addgene 52962, from Feng Zhang lab) and then infected with the lenti-Guide viral particles in presence of 8μg/ml polybrene and centrifugated for 2 h at 2900 rpm at 32°C. 48 h later, medium was replaced and Puromycin selection (10µg/mL) was applied to select positive clones for two weeks. Puromycin-resistant cells were sorted at the single cell level by FACS (Aria cell sorter). Individual clones were subjected to western blotting to confirm the absence of either *Nlrc4* or *Caspase-11* gene products. To ensure clonal reproducibility, at least 2 positive clones were compared for inflammasome response.

### Cloning and cell transduction

Galectin-3-morange construct was a kind gift from J. Enninga (Institut Pasteur, Paris, France). Galectin-3-morange coding sequence was sub-cloned into the pMSCV2.2 plasmid by first excising EGFP at the EcoRI sites followed by insertion using NotI/XhoI sites. Immortalized WT, *Nlrc4^−/−^*, *Casp1*^−/−^/*Casp11*^−/−^ or *Nlrc4*^−/−^/*Casp11*^−/−^ BMDMs were then transduced with retroviral particles, positive cells for morange fluorescence were FACS to allow clonal selection.

### Cell culture and infections

Bone-marrow derived macrophages (BMDMs) were differentiated in DMEM (Invitrogen) with 10% v/v FCS (Thermo Fisher Scientific), 10% MCSF (L929 cell supernatant), 10 mM HEPES (Invitrogen), and nonessential amino acids (Invitrogen). BMDMs were seeded in 6-, 24-, or 96-well-plates at a density of 1.25×10^6^, 2.5×10^5^, or 5×10^4^ per well. When required BMDMs were pre-stimulated overnight with 100ng/mL of PAM3CSK4 (InvivoGen). For infections with *Pseudomonas* strains, bacteria were grown overnight in Luria Broth (LB) at 37 °C with aeration. Bacterial cultures were diluted 1/50 and grew until mid-late exponential phase (OD of 1.0-1.6) and added to the macrophages at multiplicity of infection (MOI) of 25, 50 and 100, in serum and antibiotic-free medium (OPTIMEM), or as otherwise indicated. Then, to ensure homogenous infections, plates were centrifuged for 1 minute, 800 rpm. 1 h after infection, extracellular bacteria were killed by adding gentamicin (100µg/ml, Invitrogen). When specified, paraformaldehyde (PFA)-killed (PFA 4%, 20 minutes) or heat-killed (95°C, 15 minutes) of *P. aeruginosa* strains were used to infect BMDMs at MOI of 50. When required, *Shigella flexneri* (M90T), *Chromobacter violaceum* or various *Salmonella* strains (SL1344) were grown in LB in the presence or absence of antibiotics (specified in the resource table) at 37°C with constant agitation overnight. To ensure proper T3SS and flagellin expression, bacteria were sub-cultured the next day in LB media for 3 h until reaching an OD of 0,6-1.

When required, Flagellin was electroporated with Neon™ Transfection System (ThermoFischer) according manufacturer’s protocol. Briefly, 5 x 10^5^ cells were resuspended in Buffer R and 0.5 ug/ml Flagellin was electroporated in 10 μl tips using 2 pulses of 1720V and 10 width. Cells were then plated in 24 well-plates.

Transfection of cells with Fagellin (Invivogen, 0, 5µg/mL/2,5×10^5^ cells) or LPS (O111:B4, E. coli, 1µg/2,5×10^5^ cells Invivogen) was achieved using FuGeneHD (Promega) transfection reagent in Opti-MEM culture medium (44).

### Bone marrow Neutrophil isolation

Neutrophils were isolated from bone-marrow cells of WT mice by positive selection using Anti-Ly6G MicroBeads UltraPure isolation kit (Miltenyi-Biotec, Anti-Ly6G MicroBeads UltraPure mouse) according to the manufacturer’s instructions. Characterization of the purified population by Fluorescence Associated Cell Sorting showed an enrichment of more than 98% of Ly6G^high^ cells.

### Cytokine and pyroptosis measurement

IL-1β (eBioscience) was measured by ELISA. Cell-death was assayed using LDH Cytotoxicity Detection Kit (Takara). To normalize for spontaneous lysis, the percentage of LDH release was calculated as follows: (LDH infected – LDH uninfected)/(LDH total lysis – LDH uninfected)*100.

### Western blotting

Cell and supernatant protein lysates were prepared as previously described. Antibodies used were mouse anti-mouse caspase-1 antibody (Casper, Addipogen), goat anti-mouse IL-1β antibody (AF-401-NA; R&D Systems), anti-β-actin antibody (A1978; Sigma), anti-GAPDH (GTX100118; GeneTex), anti gasdermin-D (Abcam, ab209845), anti-caspase-11 (Novus, NB120-10454), anti-NLRC4 (Abcam, ab201792) and anti-NLRP3 (AdipoGen, AG-20B-0014). After incubation with primary antibody the membrane was washed 3 times with TTBS, and then Immunoreactive bands were detected by incubation for 2 h with the appropriate secondary antibodies conjugated with horseradish peroxidase (Diagomics). Proteins of interest were visualized using ECL substrate (Biorad) and images were acquired using ChemiDoc Imaging System (Biorad). Working dilutions of the antibodies are listed in **table 1**.

### Intracellular CFU experiments

2.5×10^5^ BMDMs were infected with various PAO1 strains at indicated MOIs for various times. For CFU assays after 24 hours stimulation, gentamicin protection assay was performed using gentamicin at 50µg/mL after 2 hours infections of macrophages. Supernatants were removed, and cells washed 5 times with PBS. Cells were then lysed in triton x-100 (Sigma), 0.1% and plated at 37°C overnight for CFUs numeration on LB-agar.

### Microscopy

2.5×10^5^ BMDMs were seeded on glass coverslips and infected as described above. When desired, wells were washed three times with PBS and fixed with 4% PFA for 10 minutes at 37°C. Excess of PFA was removed by PBS washes and quenching using 0,1M Glycine for 10 min at room temperature. Permeabilization and primary antibody staining were performed O/N at 4°C using Saponin 0.1%/BSA 3% solution. Stainings were performed using Hoescht (DNA labeling), rabbit anti-ASC (SCBT, 1/500), chicken anti- *P.aeruginosa* LPS (1/500, Agrisera) or anti-CD45-2-APC (1/250, Biolegends). Coverslips were washed with Saponin/BSA solution and incubated with the appropriate secondary antibodies (1/500, Diagomics). Cells were then washed 3 times with PBS and mounted on glass slides using Vectashield (Vectalabs). Coverslips were imaged using confocal Zeiss LSM 710 (Image Core facility, Biozentrum, Basel and CPTP, Toulouse) or an Olympus/Andor CSU-X1 Spinning disk microscope (Image core Facility, IPBS, Toulouse) using a 63x objective. Unless specified, for each experiment, 5-10 fields (∼50-250 cells) were manually counted using Image J software.

### Time-lapse microscopy

5×10^5^ iBMDMs Gal3-morange expressing cells were seeded in a 35 mm microdish (Ibidi). 1 hour before infection, cells were pre-stained with 0.1 µM CSFE (Biolegend) and infected with T3SS^+^ or T3SS^-^ *P. aeruginosa* with a MOI of 25 or 50. 1µM of membrane impermeant fluorescent DNA binding probe TO-PRO^®^-3 (ThermoFisher) was also added to the culture medium. Live cell imaging was performed using an Olympus/Andor CSU-X1 spinning disk microscope with a picture frequency of 5 minutes for 3-4 hours and videos were reconstructed using Image J software.

### PIT experiments

We have set up a modified form of a published protocol (13). Unless otherwise specified, WT BMDMs were infected using an MOI of 50 PAO1 T3SS^+^ or T3SS^-^ for 3 h. Intracellular bacteria-associated to PITs were visualized and manually quantified by microscopy (Olympus/Andor CSU-X1 spinning disk or inverted confocal microscope Zeiss LSM 710, ∼50 cells counted, 5-10 fields). For bacterial colony forming units snumber enumeration, cells were lysed in Triton X-100, 0.1% and plated on Luria Broth agar.

### Bacterial response to a secondary stress signal

A modified version of published protocol was used (13). WT BMDMs were infected for 2 h with PAO1 T3SS^+^ or T3SS^-^ using MOI of 50. Then, both extracellular and PIT-associated bacteria were harvested. As intracellular bacteria recovery required cell lysis with TritonX-100 0.1%, extracellular bacteria were also incubated with TritonX-100 0.1% for 2 minutes. When specified, bacteria from cells and supernatants were added to 1mL LB supplemented with 50µM Hydrogen Peroxide (H_2_0_2_) or PBS for 45 minutes, at 37°C and then plated on LB agar plates.

For neutrophil killing assays, WT BMDMs (2.5×10^5^ cells) were infected for 2 h with PAO1 T3SS^+^ or T3SS^-^ using MOI of 50. Both supernatant- and bacteria-containing PITs were exposed to neutrophils (1×10^6^ cells) for 1.5 h. Then, cells were treated with Triton X-100 0.1% for 2 minutes, and CFU counts were enumerated on LB agar plates.

### ImageStreamX experiments

WT mice (n=3 for each bacterial strain) were infected intraperitonealy with 3×10^6^ CFUs in PBS (100µL/mouse) of either T3SS^+^ or T3SS^-^ GFP-expressing bacteria. 4 hours later, peritoneal lavages were collected in 2,5mL of PBS. Neutrophils were stained prior to fixation with anti-Ly6G (APC-Vio770, Miltenyi-Biotec Clone: REA526 | Dilution: 1:50). Then, cells were fixed and permeabilized with BD Fixation/Permeabilisation Kit according to the manufacturer’s instructions, and macrophages were labeled with anti-F4/80 (BV421) (Biolegend Clone: BM8 | Dilution: 1:100). Data were acquired on ImageStreamX^MKII^ (Amnis) device (CPTP Imaging and Cytometry core facility) and analyzed using IDEAS software v2.6 (Amnis). The gating strategy used to evaluate efferocytosis of bacteria-containing macrophages by Neutrophils was performed as follows: (i) gate was set on cells in focus and (ii) sub-gated on single cells. Then, we gated both on (iii) Ly6G^+^ Neutrophils and on (iv) F4/80^+^ macrophages within Ly6G^+^ population. (v) To distinguish efferocytosis of intact from fragmented macrophages, we created a mask based on the Surface Area of F4/80+ signal. This was applied to Ly6G^+^/F4/80^+^ gate. (vi) The percentage of GFP^+^ bacteria within the Ly6G^+^/F4/80^+^ population was visualized and quantified.

### Bacterial KO generation and complementation

The knockout vector pEXG2 was constructed and used based on the protocol described by Rietsch et al. (45) with the following modifications. Briefly, 700-bp sequences of the flanking regions of the selected gene were amplified by PCR with Q5 high fidelity polymerase (New England Biolabs). Then, the flanking regions were gel purified and inserted into pEXG2 plasmid by Gibson assembly (46). The assembled plasmid was directly transformed into competent SM10λpir using Mix&Go competent cells (Zymo Research Corporation) and plated on selective LB plates containing 50 µg/mL kanamycin and 15 µg/mL gentamicin. The resulting clones were sequenced, and mating was allowed for 4 h with PAO1 strain at 37°C. The mated strains were selected for single cross over on plates containing 15 µg/mL gentamicin and 20 µg/mL Irgasan (removal of *E.coli* SM10 strains). The next day, some clones were grown in LB for 4 hours and streaked on 5% sucrose LB plates overnight at 30°C. *P. aeruginosa* clones were then checked by PCR for mutations. For flagellin complementation experiments, JBOC plasmid (homemade) was used to re-express flagellin gene (*fliC*) under its endogenous promoter in *fliC* deficient PAO1 strains. Briefly, PAO1 *fliC* and its promoter were PCR-amplified, purified and integrated in JBOC plasmid. SM10 *E.coli* were transformed with *fliC* plasmid and allowed to conjugate with PAO1 mutant strains as described above. Positive PAO1 strains were selected on Gentamicin-Irgasan plates and checked by subsequent PCRs. All primers were designed with Snapgene software (GSL Biotech LLC).

### Statistical analysis

Statistical data analysis was performed using Prism 5.0a (GraphPad Software, Inc.). For comparison of two groups (cell death, cytokine secretion, CFU and microscopy-based counts), we used two-tailed t-test. Bonferroni correction was applied for multiple comparisons to two-way ANOVA statistical analysis. Data are reported as mean with standard deviation (s.d). For animal experiments Mann-Whitney tests were performed. P values are given in figures, NS means non-significant, unless otherwise specified. Significance is specified as *, ** or *** for P-values <0.05, <0.01 or <0.001 respectively.

### Data and software

Fiji and Image J software was used for time-lapse and microscopy images processing and analysis. Snapgene software (GSL Biotech LLC) was used to design all primers. Cytometry data analysis and processing were performed using FlowJo.10 software. ImageStreamX data were analyzed using IDEAS software v2.6 (Amnis). Benchling online software was used to design Crispr guides.

### Ethics Statement

All animal experiments were approved (License 2535, Kantonales Veterinäramt Basel-Stadt and License APAFIS#8521-2017041008135771, Minister of Research, France) and performed according to local guidelines (Tierschutz-Verordnung, Basel-Stadt and French ethical laws) and the European Union animal protection directive (Directive 2010/63/EU).

## Acknowledgements

We thank the animal facilities, image core facilities and FACS core facility from Biozentrum (Univ. of Basel, Switzerland), IPBS (CNRS, Univ. of Toulouse, France) and CPTP (INSERM, Univ. of Toulouse, France) institutes. Specifically, we acknowledge (S. Allard and D. Daviaud from the CPTP imaging facility. Authors acknowledge V.M Dixit (Genentech) (*Nlrc4*^−/−^, *Casp11*^−/−^), R. Flavell (*Casp1*^−/−^*11*^−/−^), F. Sutterwala (*Nlrc4*^−/−^) for sharing mice. We also acknowledge J. Enninga (Institut Pasteur, Paris, France) for galectin-3 plasmid share and R.E. Vance and I. Rauch (University of California at Berkeley) for providing bones from mice deficient for *Naip5*^−/−^ and *Naip*^Δ1-6^. We also thank Y. Rombouts and G.L. Villarino for manuscript reading and comments. We also thank funding partners. E.E. and E.M. are funded by a FRM “Amorçage Jeunes Equipes” number AJE20151034460. E.M. is also funded by an ATIP-AVENIR grant and an ERC StG INFLAME 804249. P.B. is funded by the Swiss National Science Foundation (310030_165893). We thank Life Science Editors for editing assistance. Authors declare no conflict of interest.

## Author Contribution

E.E, R.P, J.B, P.B and E.M designed the study; D.B, A.Coste and O.N provided essential mouse lines, reagents and expertize. E.E, R.P, J.B, P.J.B, A.Colom, O.C, R.F.D, J.V.S, C.C and E.M performed experiments; E.M wrote the manuscript, with contribution from O.N.

## Supplementary information

**Figure S1:**
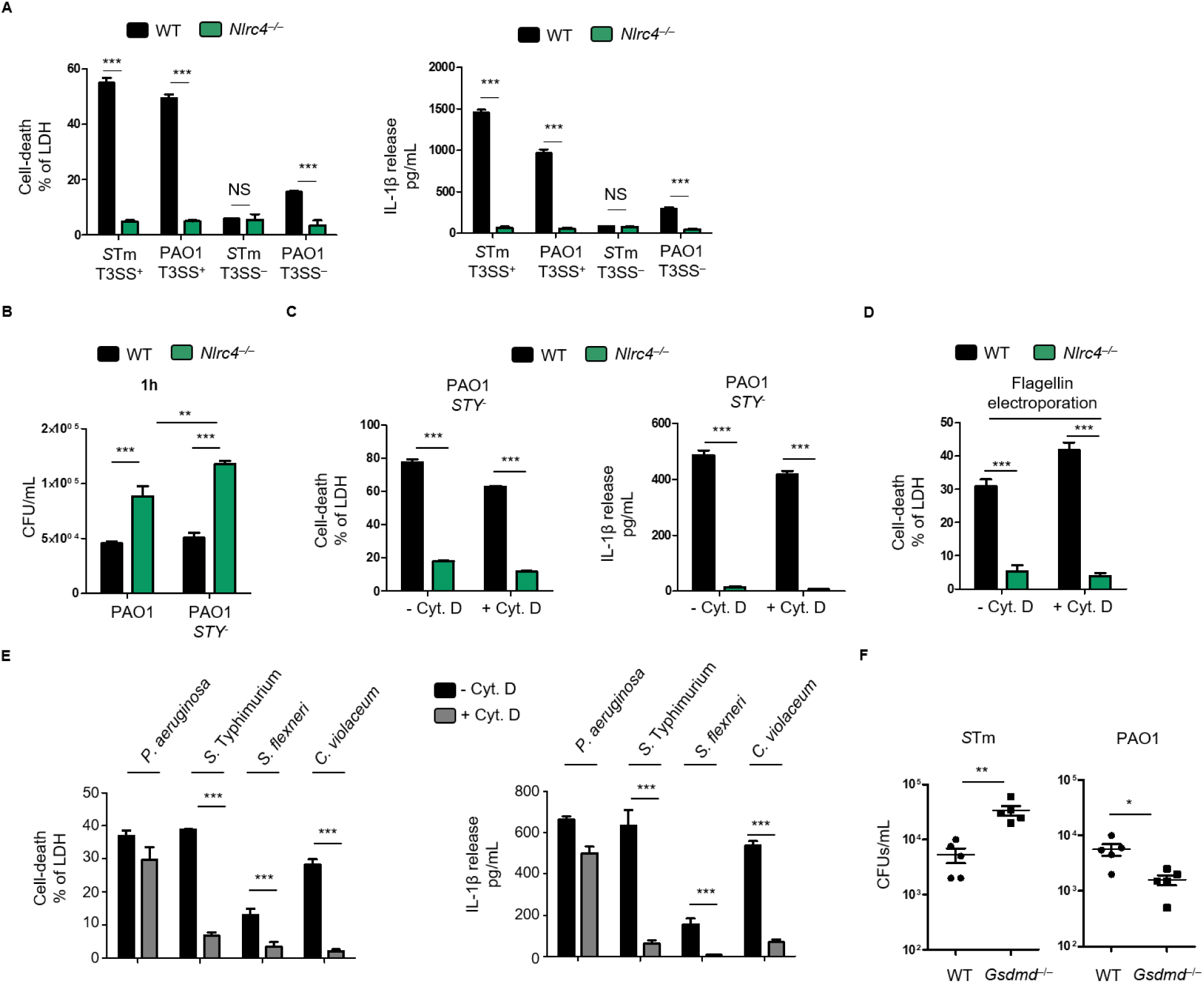
*Pseudomonas aeruginosa* T3SS promotes phagocytosis-independent activation of the NLRC4 inflammasome. BMDMs were primed with 100 ng/mL of the TLR2 ligand Pam_3_CSK_4_ for 16 h to induce pro-IL-1β expression and then infected for 3 hours (h) with various bacteria (MOI 15) or inflammasome inducers. **(A)** Measurement of cell death (LDH release) from WT and *Nlrc4^−/−^* BMDMs after 3 h of infection with either T3SS^+^ or T3SS^-^ PAO1 or *S*Tm. **(B)** CFU scoring in WT and *Nlrc4^−/−^* BMDMs infected for 1h with either PAO1 or *STY*- deficient *P. aeruginosa*. **(C)** LDH and IL-1β release quantifications of WT and *Nlrc4*^−/−^ BMDMs infected for 3 h with either PAO1 or *STY*-deficient *P. aeruginosa* in presence or not of Cytochalasin D (0,2 µg/mL). **(D)** LDH release quantifications of WT and *Nlrc4*^−/−^ BMDMs, pretreated or not with Cytochalasin D (0,2 µg/mL), electropororated (Neon™ Transfection System) with 0.5 ug/ml and then plated for 1 h. **(E)** LDH and IL-1β released by WT BMDMs, pre-incubated or not with cytochalasin D (0.2µg/mL) for 30 minutes, and then infected with either *P. aeruginosa*, *S*. Typhimurium, *S. flexneri* or *C. violaceum*. **(F)** CFU scoring in the peritoneal cavity of WT and *GsdmD*^−/−^ mice infected for 6 h with 3.10^6^ CFUs of *P. aeruginosa* and *S*. Typhimurium. All graphs **(A-E)** show mean and s.d of quadruplicate wells from three pooled experiments. **(F)** represents one experiment from three independent experiments.

**Figure S2:**
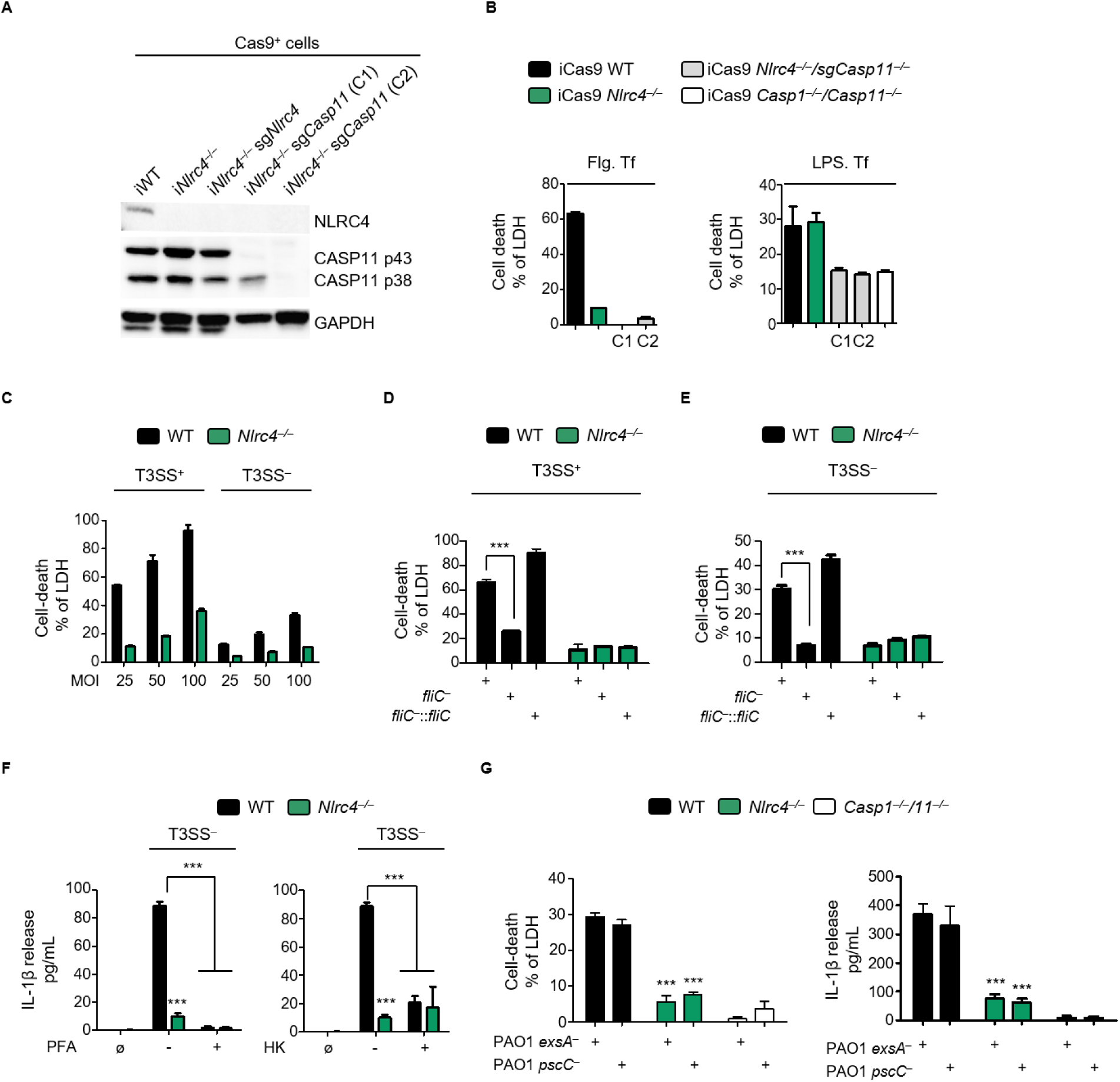
*P. aeruginosa* triggers T3SS-independent activation of both NLRC4 and Caspase11 inflammasomes. **(A, B)** Biochemical **(A)** and functional **(B)** characterization of two iCas9*Nlrc4*^−/−^/sg*Casp11*^−/−^ CRISPR KO clones. **(B)**, LDH release measurement of CRISPR clones (A) transfected with either 0,5 µg/mL of Flagellin (2 hours) or 5µg/mL of LPS (10 hours). **(C)** Cell death evaluation (LDH) in WT and *Nlrc4*^−/−^ BMDMs infected with various MOIs of PAO1 T3SS^+^ or T3SS^-^ for 3 h. **(D, E)** LDH measurement in WT and *Nlrc4*^−/−^ BMDMs infected by PAO1 T3SS^+^, T3SS^-^ or T3SS^-^/*fliC*^-^, complemented or not for flagellin expression (*fliC^-^*:: *fliC*) for 3h with an MOI of 50. **(F)** Measurement of IL-1β release from WT and *Nlrc4^−/−^* BMDMs after 3 h of infection with live, PFA-killed or heat killed (HK) T3SS deficient-PAO1 at an MOI of 50. **(G)** Measurement of LDH and IL-1β release from WT, *Nlrc4*^−/−^ and *Casp1*^−/−^*/11*^−/−^ BMDMs after 3 h of infection with T3SS deficient-PAO1 *exsA*^-^ or -PAO1 *pscC*^-^ at an MOI of 50. Graphs show mean and s.d of quadruplicate wells pooled from three independent experiments.

**Figure S3:**
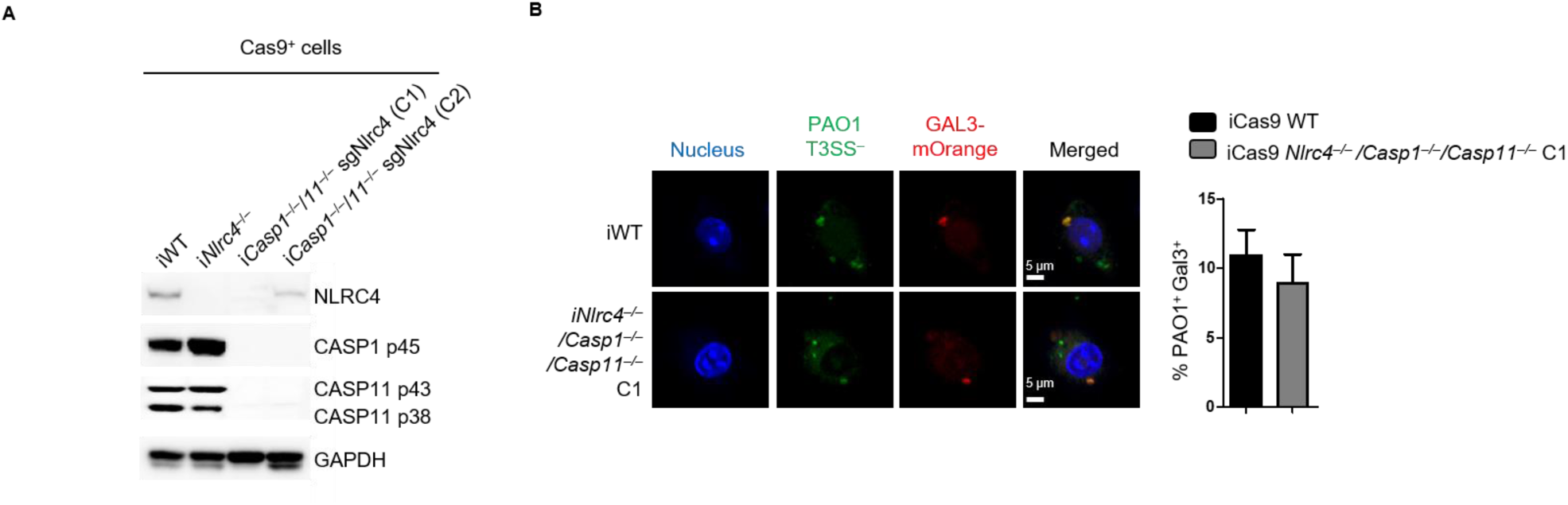
T3SS- *P. aeruginosa* associate to altered phagosomes. **(A)** Biochemical characterization of two iCas9*Nlrc4*^−/−^/sg*Casp11*^−/−^ CRISPR KO clones. **(B)** Fluorescence microscopy observations and quantifications of galectin-3 targeted PAO1 T3SS^-^ (MOI 25, 3 h of infection) in iWT and i*Casp1^−/−^/11^−/−^*/sg*Nlrc4*^−/−^ BMDMs transduced with galectin-3-morange. Graphs show mean and s.d of quadruplicate wells from three independent pooled experiments. Immunoblotting representative of one experiment performed on various clones. Microscopy analysis visualized 10 fields containing approximately 200 cells using Image J software.

**Figure S4:**
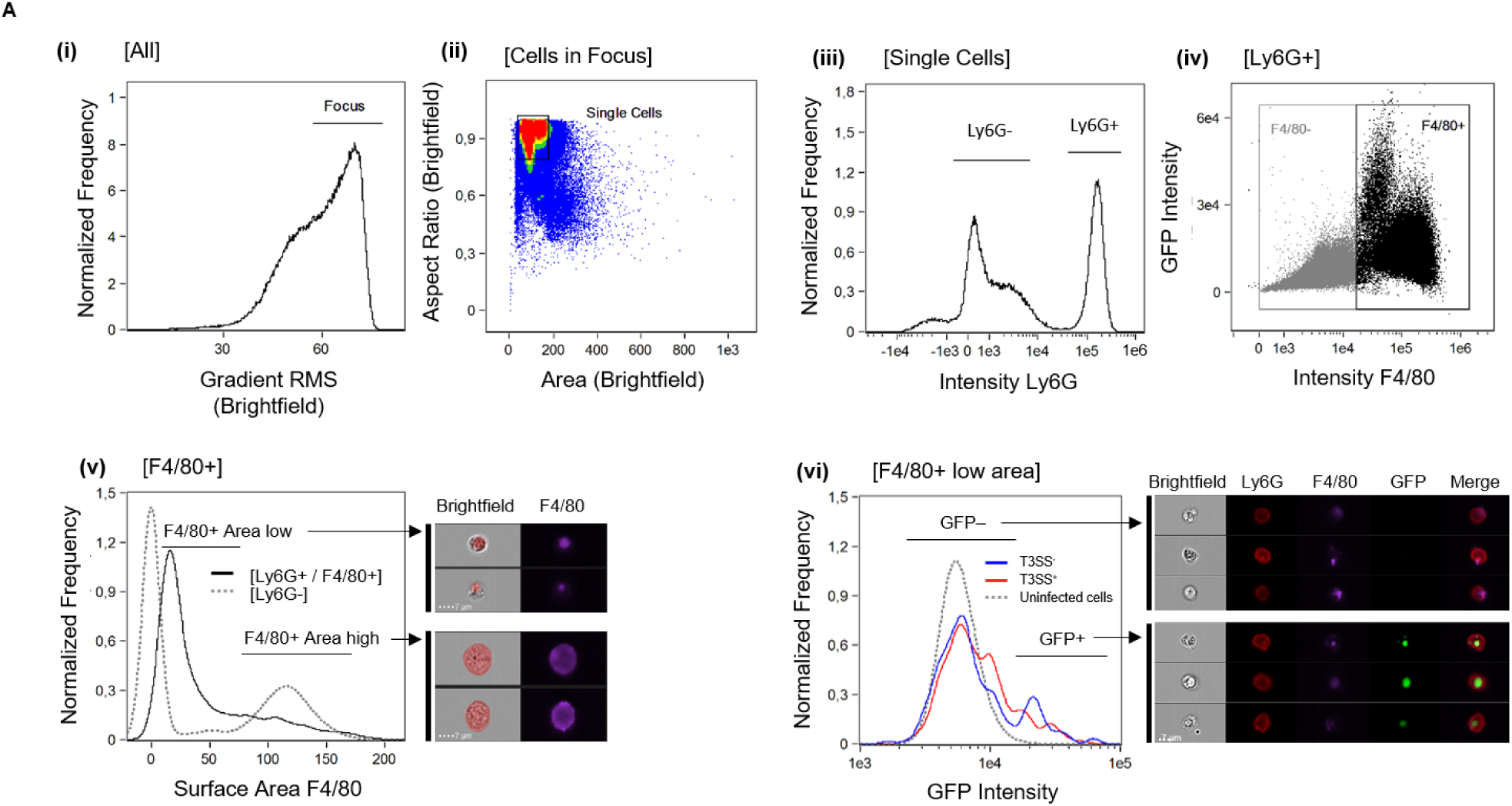
ImagestreamX gating strategy used. **(A)** ImagestreamX gating strategy used to visualize and quantify neutrophil-mediated efferocytosis of bacteria-containing macrophages. (i) A gate was set on cells in focus [Cells in Focus], then (ii) single cells [Single Cells] were gated. (iii) First, gate was put on Ly6G^+^ Neutrophils [Ly6G^+^] and then we selected (iv) F4/80^+^ macrophages [F4/80^+^] within Ly6G^+^ population. (v) A mask based on the surface Area of F4/80^+^ signal was applied to Ly6G^+^/F4/80^+^ gate to discriminate efferocytosis of intact macrophages from fragmented macrophages. As a control, dotted gray line shows surface area of F4/80^+^ signal in the [Ly6G^-^] gate. Then, (vi) we gated on [F4/80^+^ low Area] and measured the intensity of the GFP signal. Red and Blue lines show the intensity of GFP signal measured in mice challenged with either *P. aeruginosa* T3SS^+^ or T3SS^-^ strains respectively. Finally, GFP^+^ bacteria percentage within the gate (vi) was visualized. Upper image panel and lower image panel show representative images from the Neutrophils (Ly6G^+^)/macrophages (F4/80^+^) gate obtained in GFP^-^ gate or GFP^+^ gate respectively.

**Figure S5:**
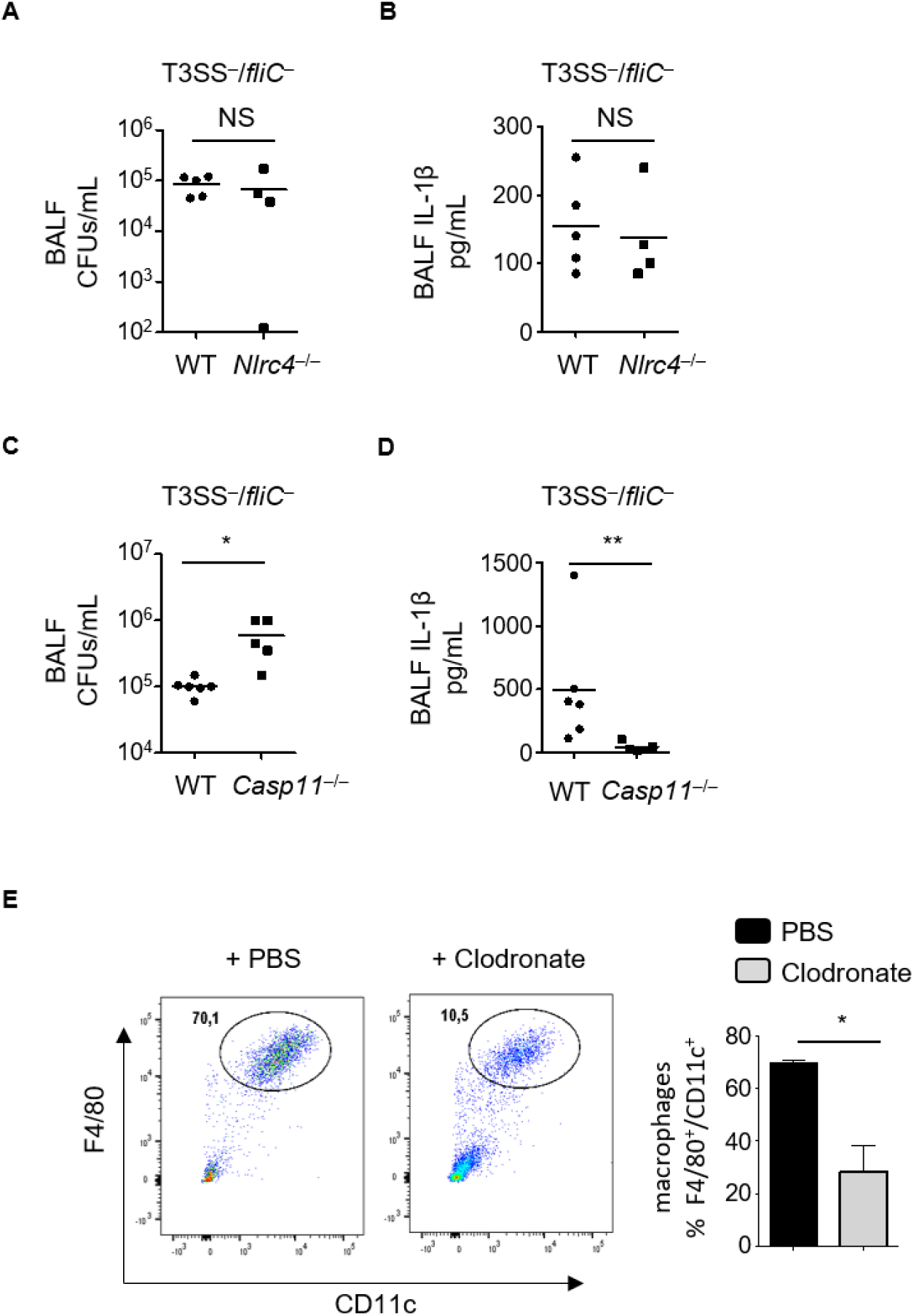
NLRC4 and Caspase-11 only protect mice against T3SS-deficient *P. aeruginosa*. **(A-D)** PAO1 CFUs scoring and IL-1β levels in BALFs from WT, *Nlrc4*^−/−^ **(A, B)** or *Casp11*^−/−^ **(C, D)** mice after 18 h of infection with T3SS^-^/*fliC*^-^ *P. aeruginosa* strain with 1.5×10^7^ CFUs. **(E)** FACS representation and quantification of depleted alveolar macrophages (F4/F80^+^, CD11c^+^) in control (PBS) and clodronate instilled mice.

**Table S1:**
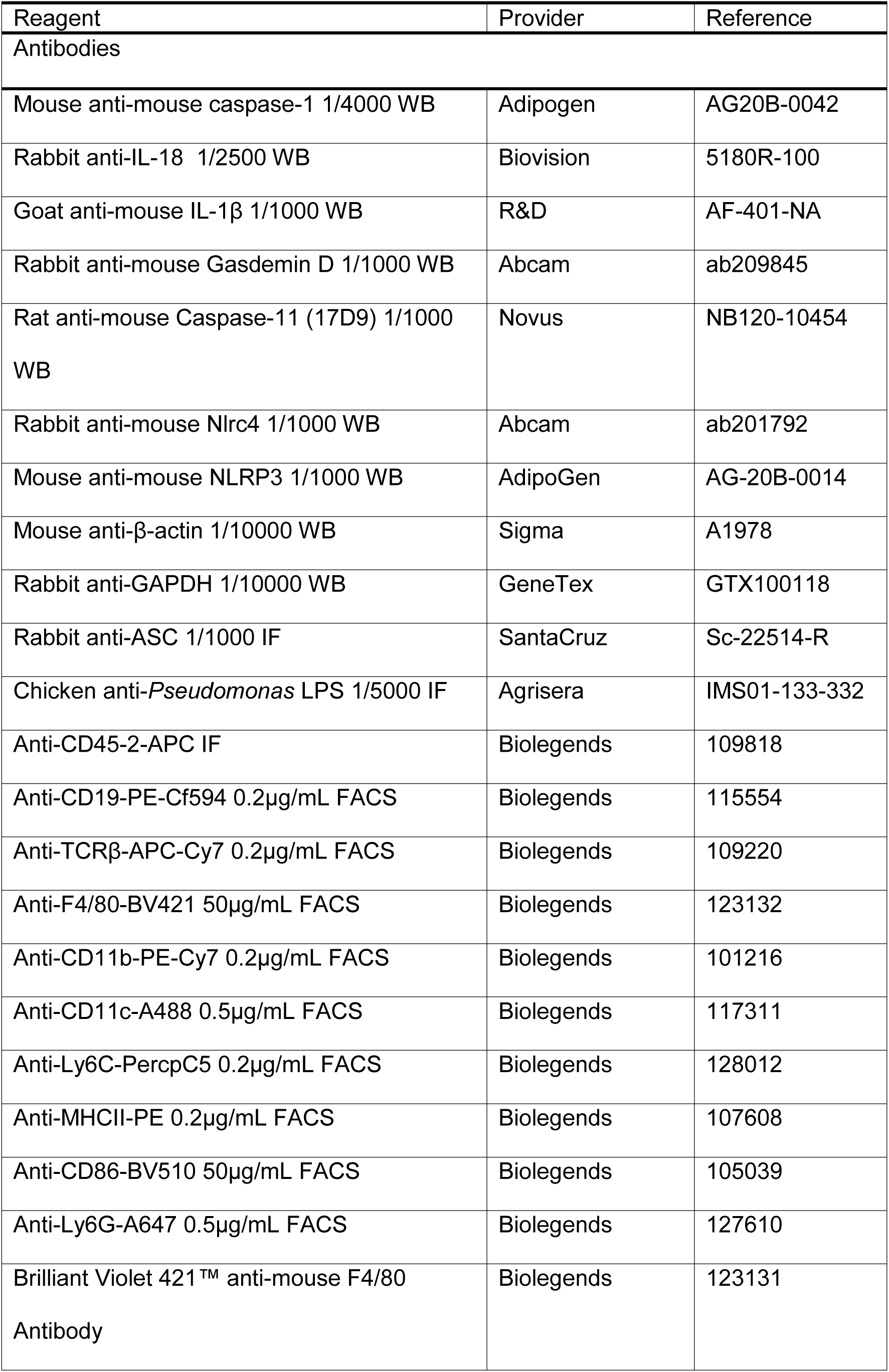
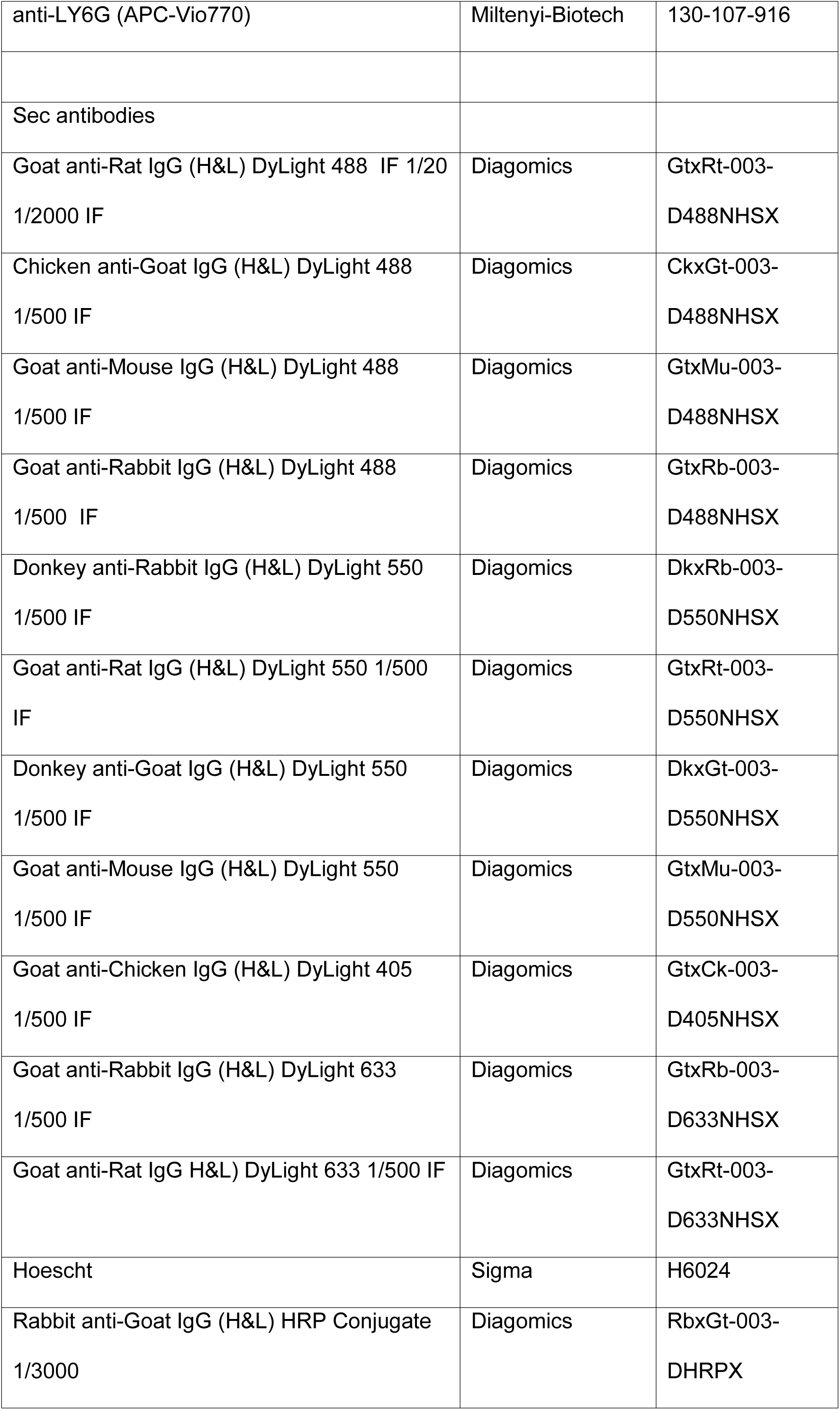
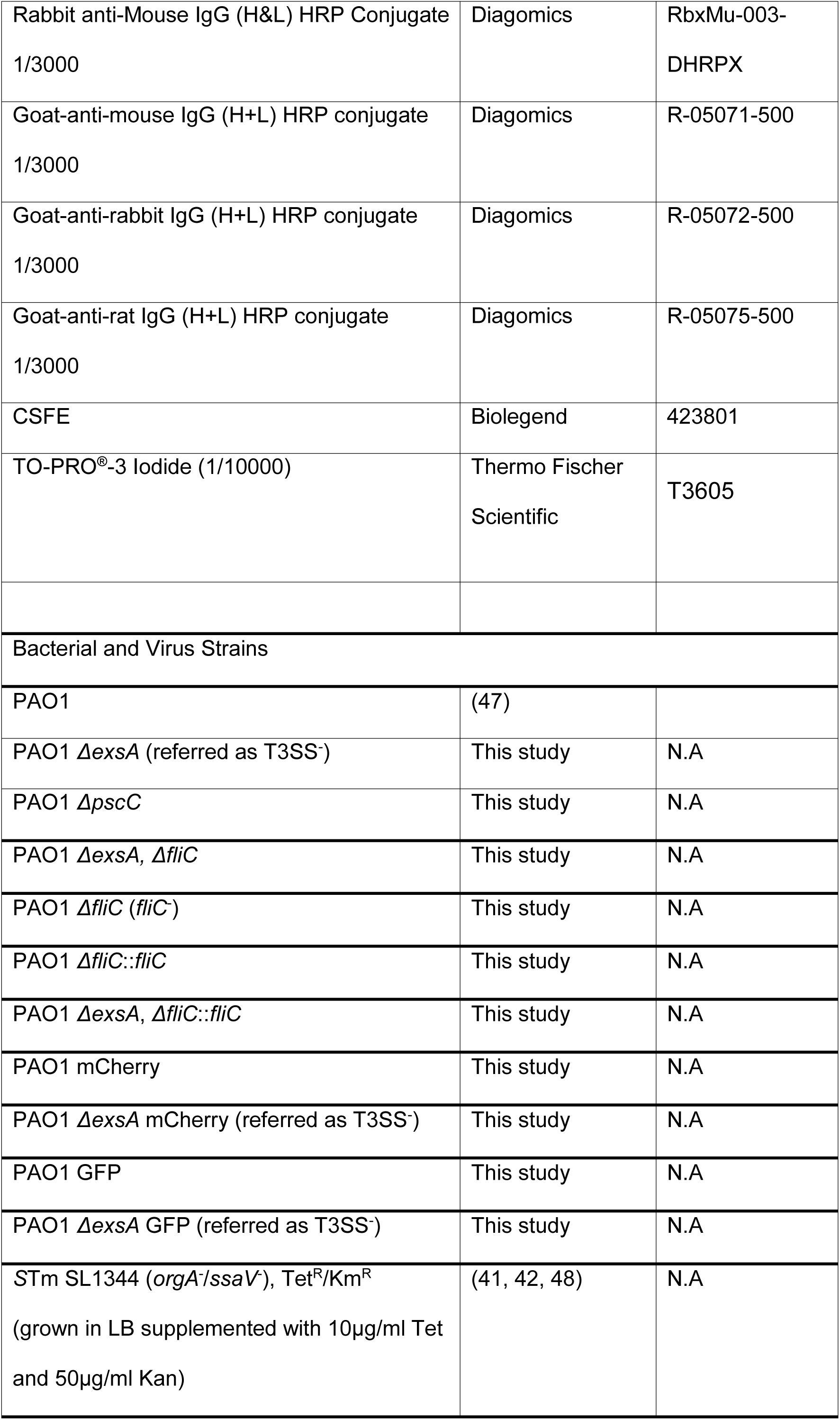
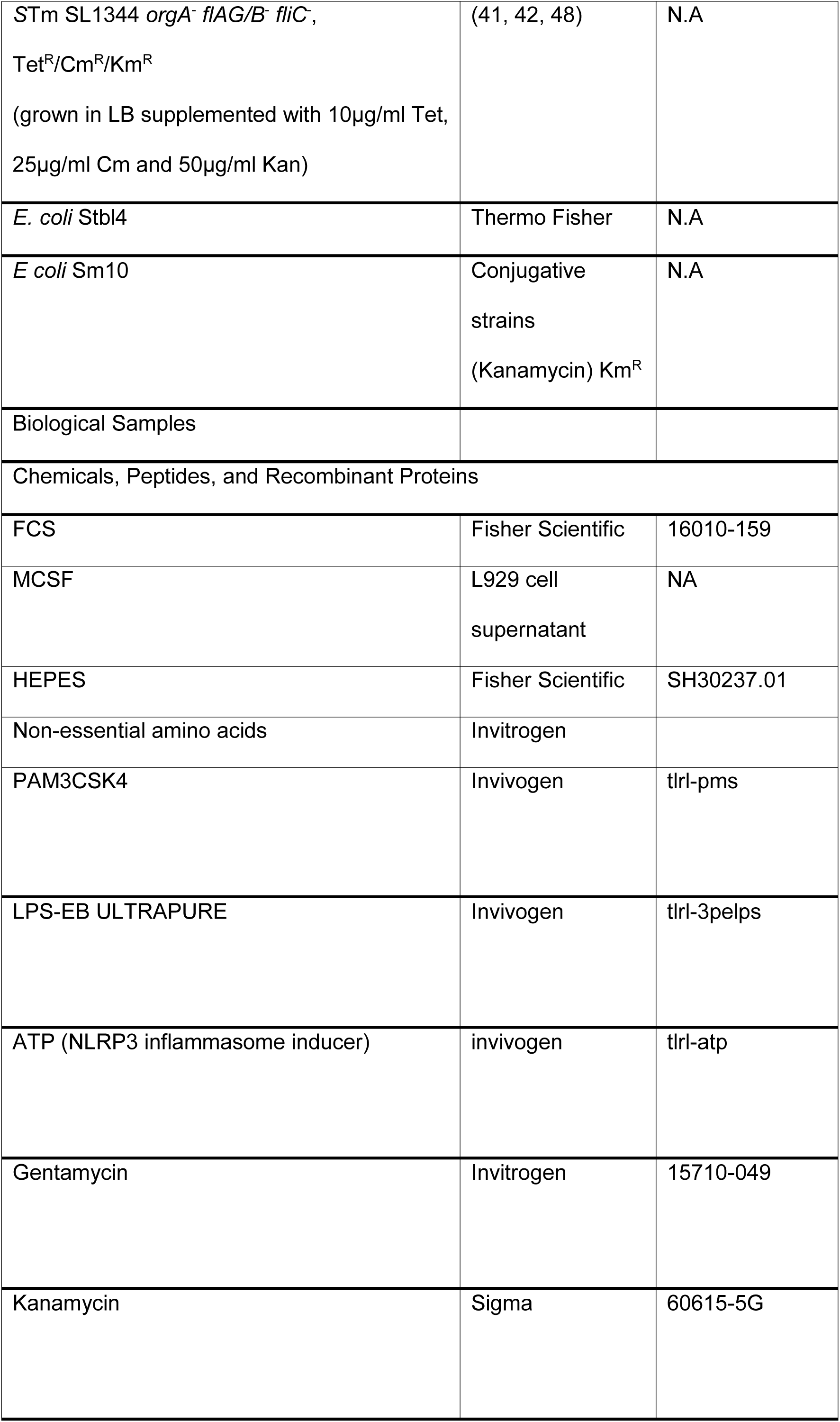
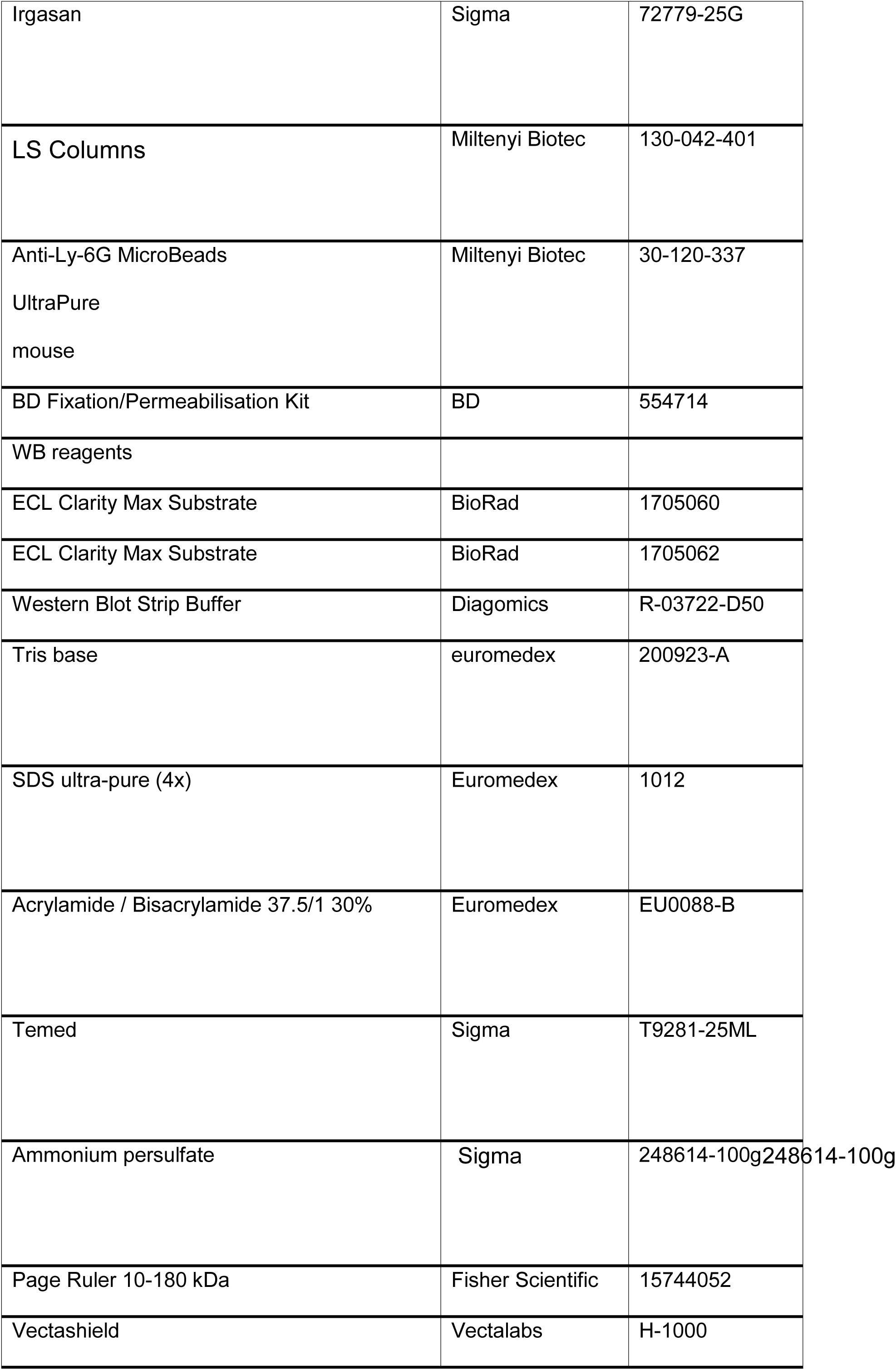
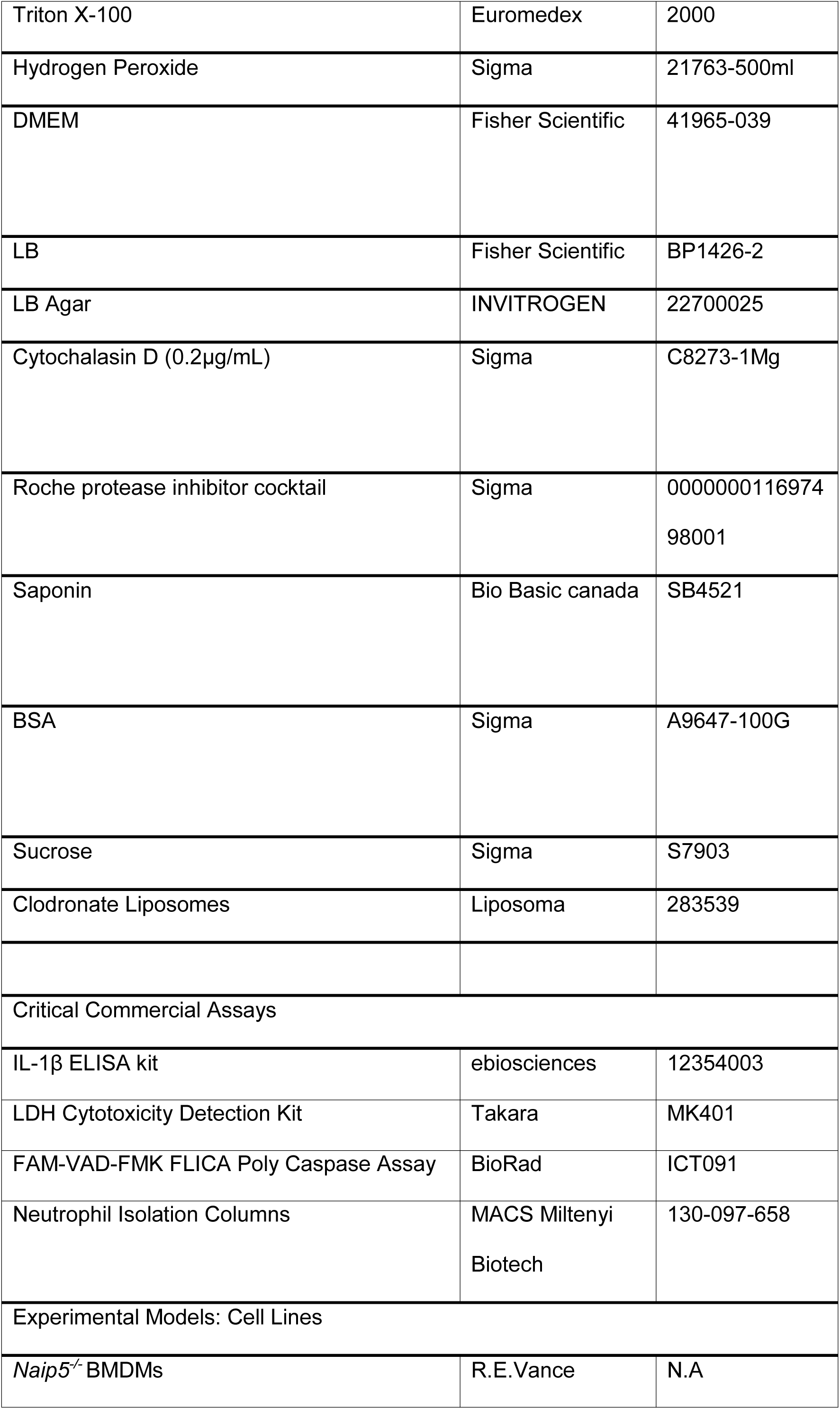
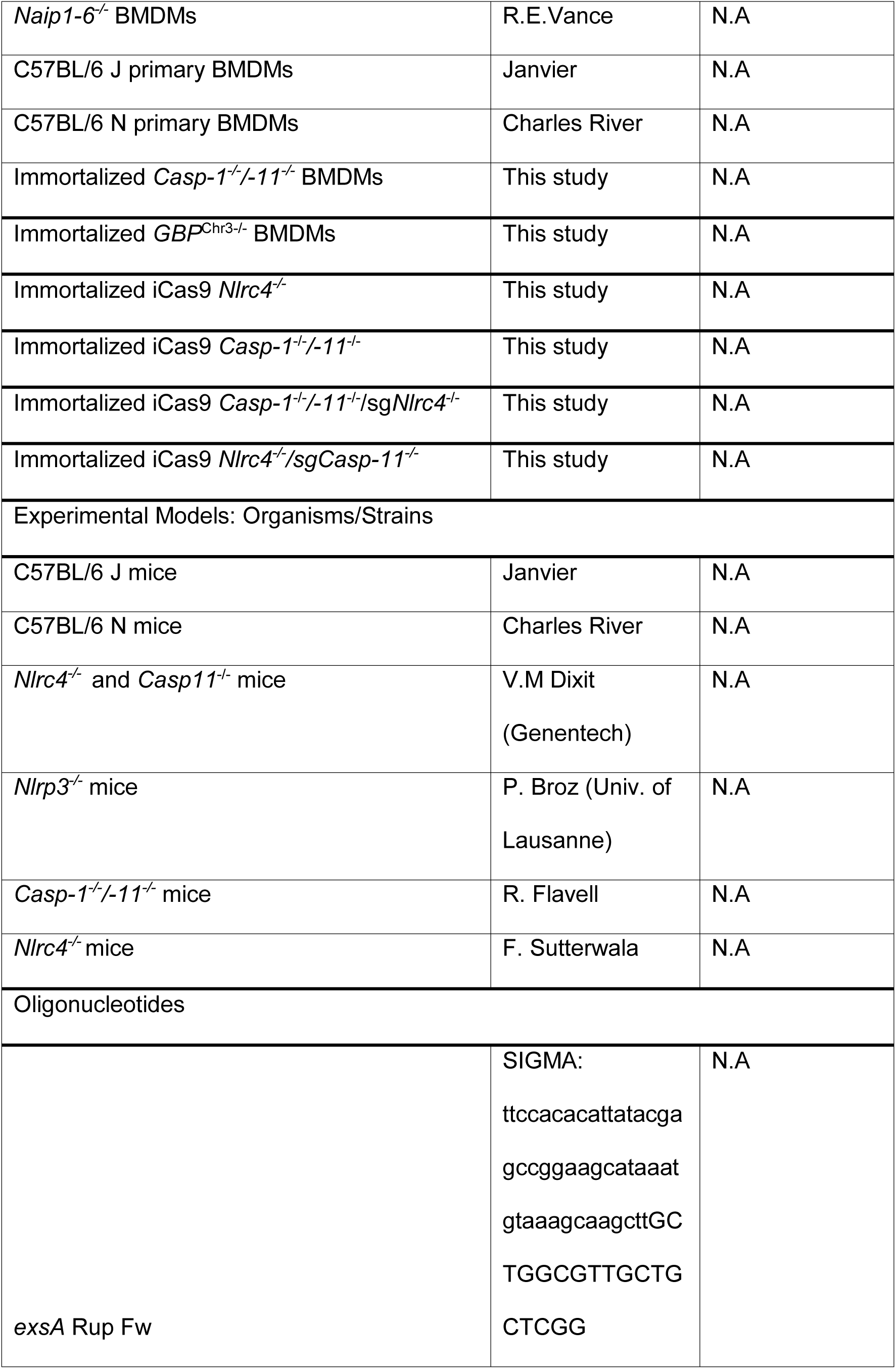
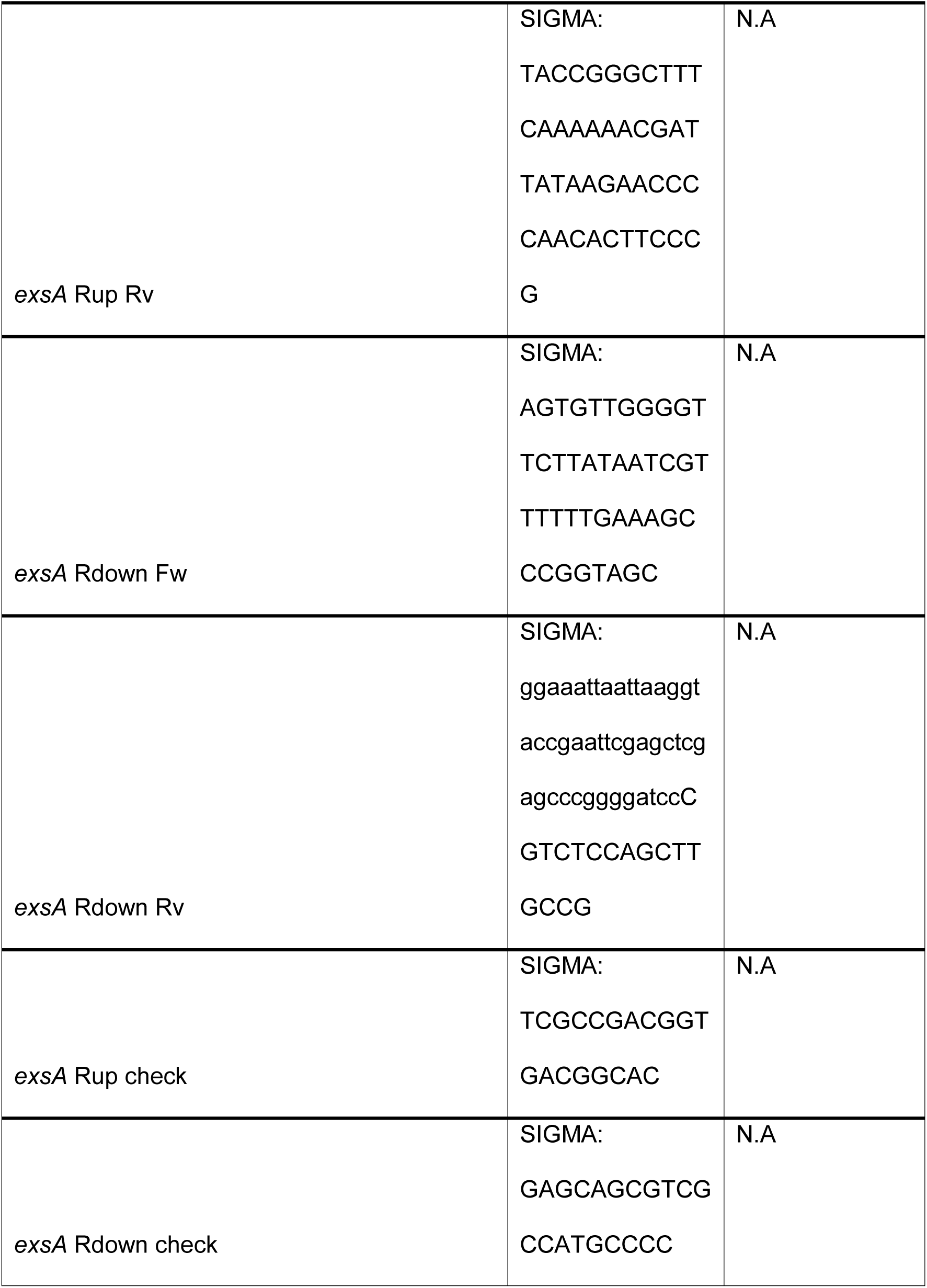
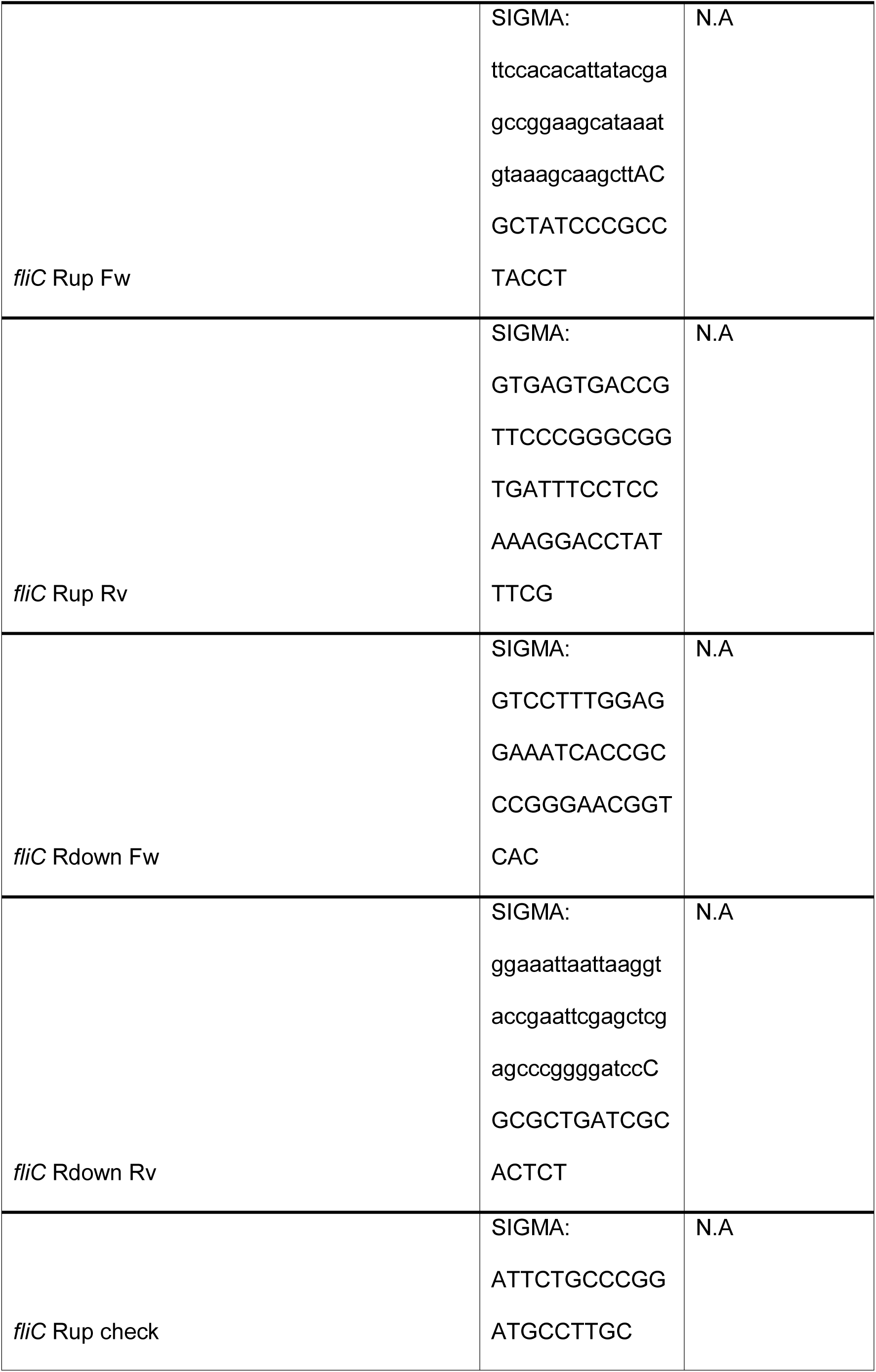
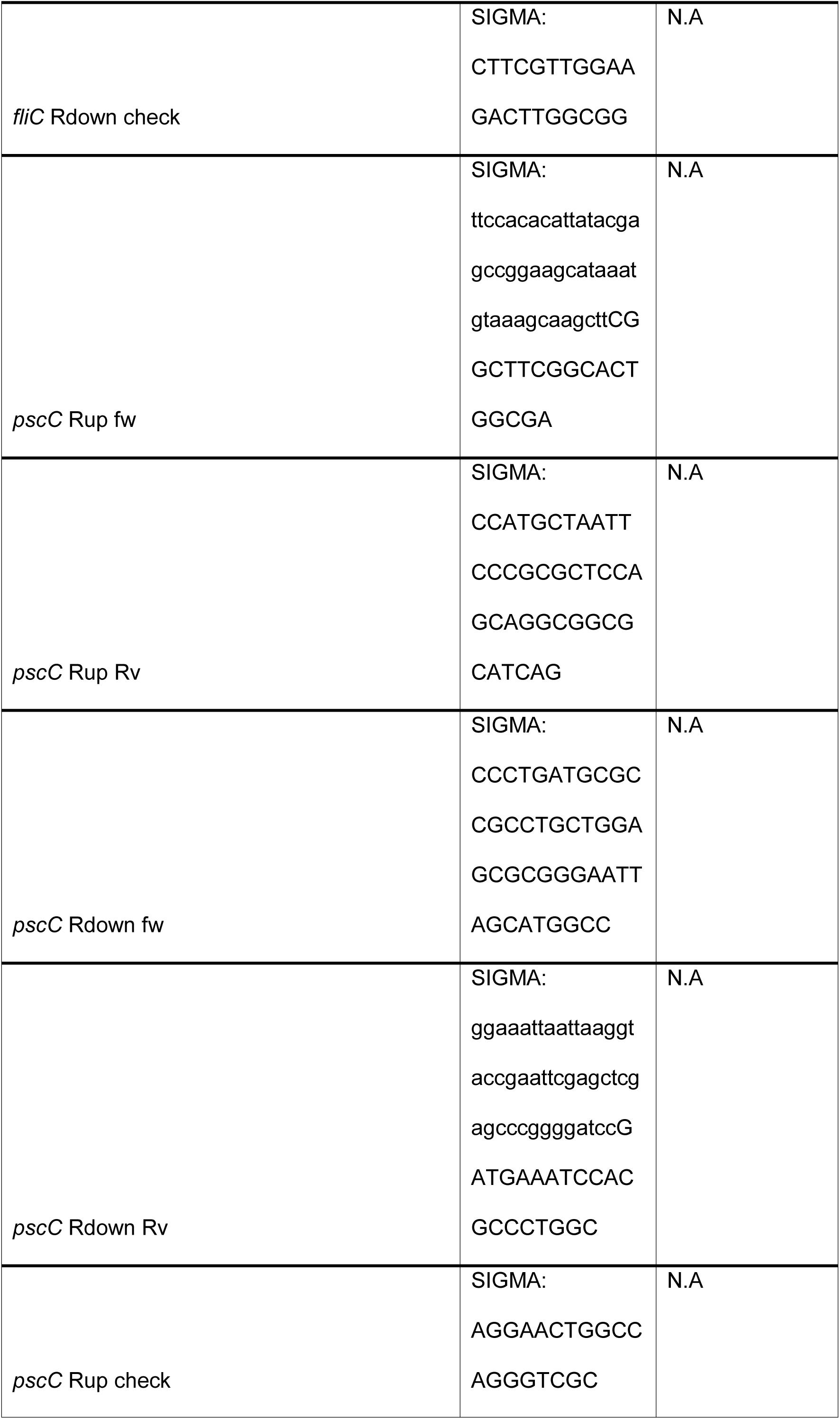
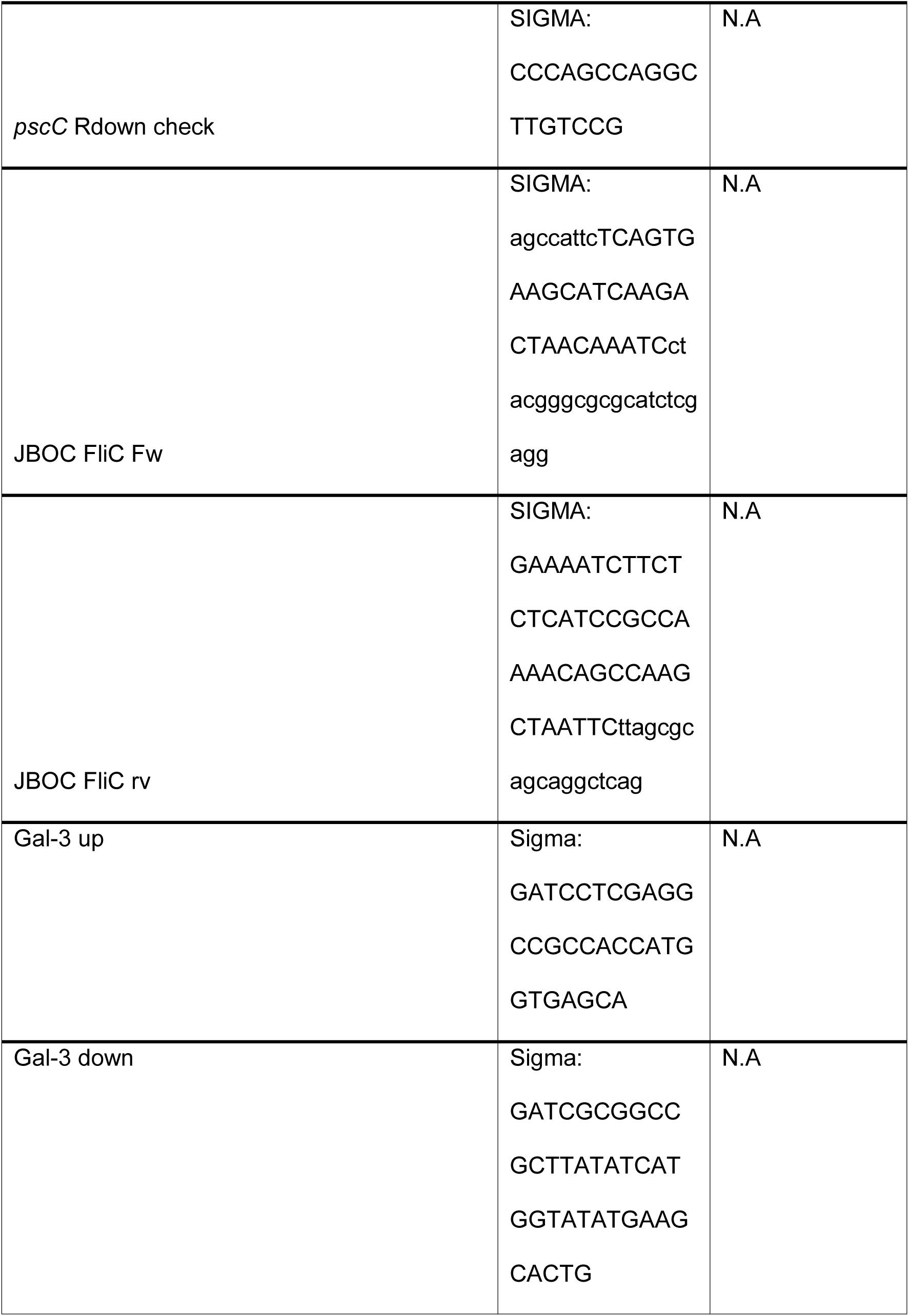
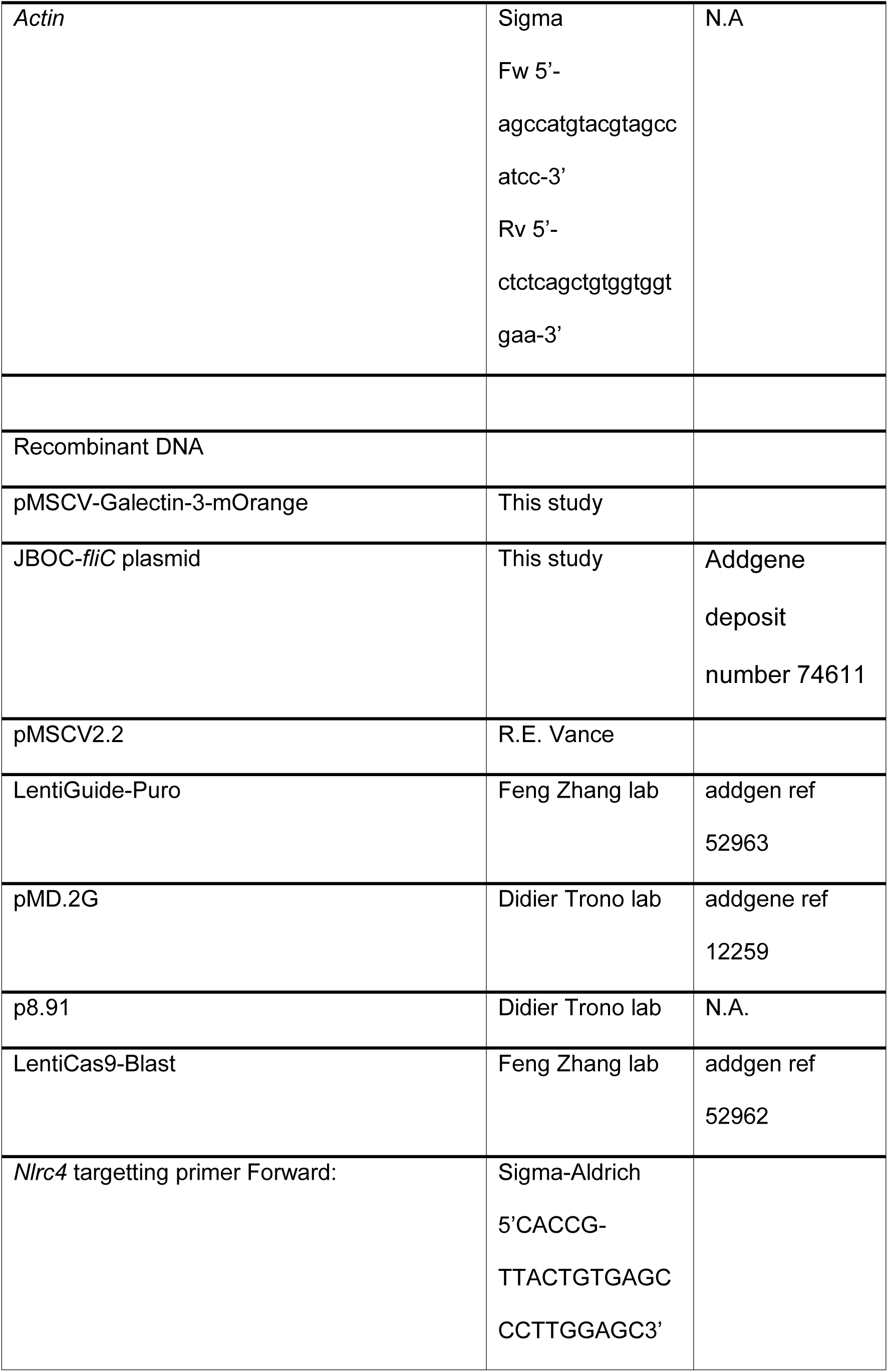
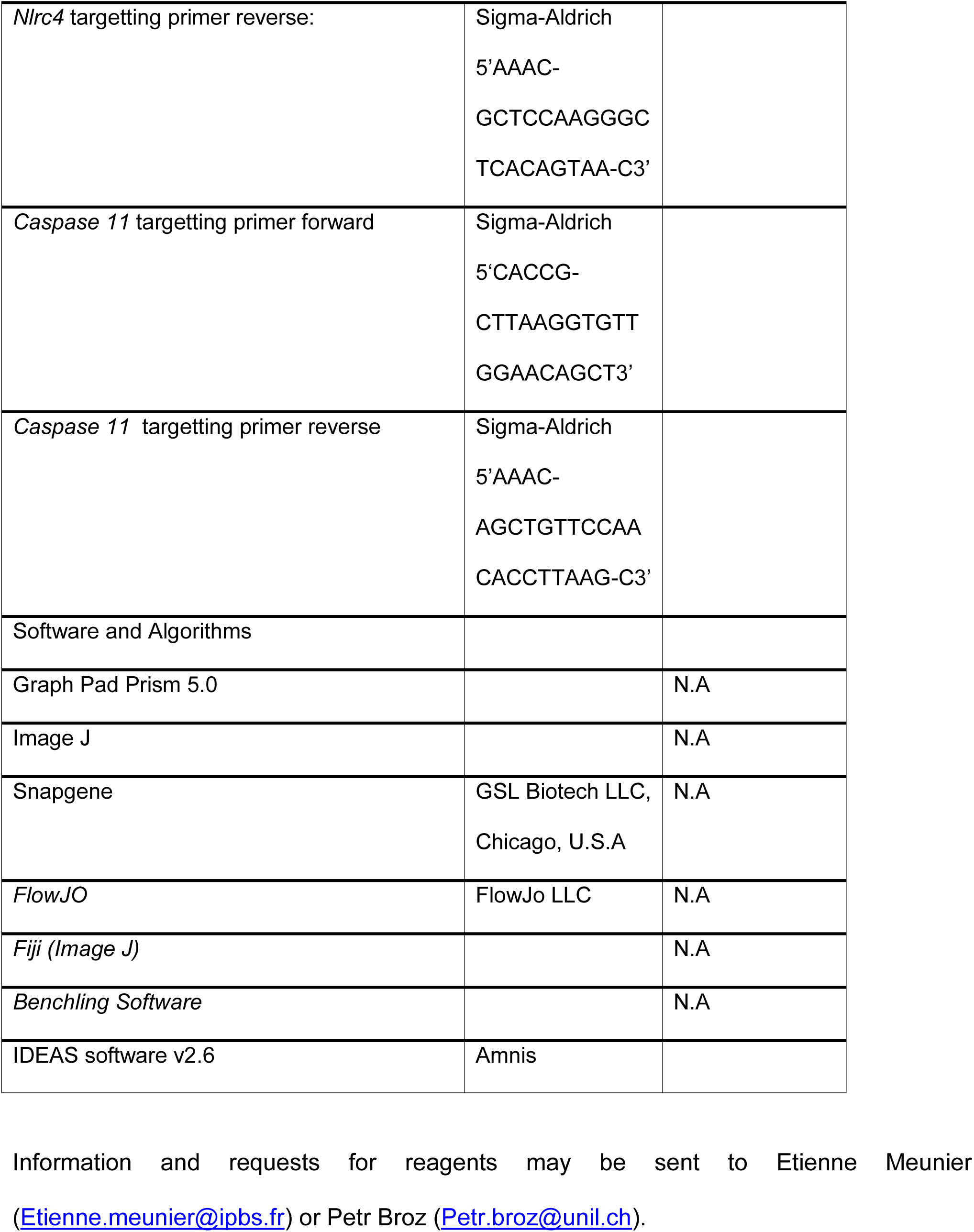
All products, software and biological samples used in the current study, including their references and concentrations, are listed in table 1.

